# Accelerated ensemble generation for cyclic peptides using a Reservoir-REMD implementation in GROMACS

**DOI:** 10.1101/2022.09.07.507010

**Authors:** Shawn C.C. Hsueh, Adekunle Aina, Steven S. Plotkin

## Abstract

Cyclic peptides naturally occur as antibiotics, fungicides, and immunosuppressants, and have been adapted for use as potential therapeutics. Scaffolded cyclic peptide antigens have many protein characteristics such as reduced toxicity, increased stability over linear peptides, and conformational selectivity, but with fewer amino acids than whole proteins. The profile of shapes presented by a cyclic peptide modulates its therapeutic efficacy, and is represented by the ensemble of its sampled conformations. Although some algorithms excel in creating a diverse ensemble of cyclic peptide conformations, they seldom address the entropic contribution of flexible conformations, and they often have significant practical difficulty producing an ensemble with converged and reliable thermodynamic properties. In this study, an accelerated molecular dynamics (MD) method, reservoir replica exchange MD (R-REMD or Res-REMD), was implemented in GROMACS-4.6.7, and benchmarked on three small cyclic peptide model systems: a cyclized segment of A*β* (cyclo-(CGHHQKLVG)), a cyclized furin cleavage site of SARS-CoV-2 spike (cyclo-(CGPRRARSG)), and oxytocin (disulfide bonded CY-IQNCPLG). Additionally, we also benchmarked Res-REMD on Alanine dipeptide and Trpzip2 to demonstrate its validity and efficiency over REMD. Compared to REMD, Res-REMD significantly accelerated the ensemble generation of cyclo-(CGHHQKLVG), but not cyclo-(CGPRRARSG) or oxytocin. This difference is due to the longer auto-correlation time of torsional angles in cyclo-(CGHHQKLVG) *v* s. the latter two cyclic peptide systems; The randomly seeded reservoir in Res-REMD thus accelerates sampling and convergence. The auto-correlation time of the torsional angles can thus be used to determine whether Res-REMD is preferable to REMD for cyclic peptides. We provide a github page with modified GROMACS source code for running Res-REMD at https://github.com/PlotkinLab/Reservoir-REMD.

## 1 Introduction

Structural ensemble generation has become an important tool to sample and characterize the underlying energy landscape governing the kinetics, thermodynamics, and functionally important motions of biomolecules, ^1–5^ an area of research of fundamental importance, in which José Onuchic has had a significant role in establishing the theoretical and computational paradigms guiding the field. ^6–21^ A molecular dynamics application of parallel tempering,^22^ Replica Exchange Molecular Dynamics (REMD), ^23–25^ has been widely used to facilitate accurate sampling of the energy landscape^26,27^ REMD is a more powerful tool than conventional molecular dynamics simulations for energy landscape sampling, because conformations in high-temperature replicas can easily overcome energy barriers, and the diverse structures sampled in high-temperature replicas can traverse to the lower-temperature replica of interest to accelerate sampling. However, REMD is also computationally expensive. The ensemble generated by low-temperature replicas cannot converge until the high-temperature replica samples all the representative stable states. This “burn-in” process can be avoided by separating REMD into two steps: The initial generation of stable representative structures at the highest-temperature replica, and then the ensemble generation at the other lower temperature replicas. A modified REMD method, Reservoir Replica Exchange Molecular Dynamics (R-REMD),^28^ utilizes a pre-sampled high-temperature reservoir to seed the REMD simulation, wherein this reservoir is coupled to the multiparallel REMD simulation through exchange attempts between a *randomly drawn state* from the reservoir and the highest temperature replica of the REMD simulation (which is at slightly lower temperature than the reservoir). While the written form of R-REMD is unambiguous, we have found the acronym R-REMD awkward to articulate and conducive to verbal miscommunication; We have adopted “Res-REMD” in our discussions and will continue to use that convention here.

Prior application of R-REMD/Res-REMD has successfully yielded conformational ensembles in good agreement with conventional REMD for a wide variety of small molecules and peptides such as butane,^29^ leucine dipeptide in gas phase, ^29,30^ leucine tripeptide in implicit solvent,^30^ Trpzip2 and DPDP in implicit solvent,^28^ A*β*_21*−*30_ peptide in explicit solvent, ^31^ vasopressin and oxytocin and their mutants in explicit solvent,^32^ and also for dithiacyclophane and cinchona alkaloid organocatalyst using ab initio energy functions. ^33^ A variant of Res-REMD, non-Boltzmann R-REMD, also yielded conformational ensembles in good agreement with conventional REMD for alanine tetrapeptide, alanine undecapeptide, and Trp-cage miniprotein in implicit solvent,^34^ Trpzip2 in implicit solvent,^35^ Alanine dipeptide, and RNA (rGACC) tetramer in explicit solvent.^36^ Res-REMD has only been available in AMBER software^37^ however, and only recently made available in Python.^33^ In this study, Res-REMD is implemented in GROMACS-4.6.7 for the first time, and benchmarked on several cyclic peptides as well as two other common model systems, Alanine dipeptide and Trpzip^2^.

Cyclic peptides have been widely used as potential therapeutics^38–40^ and scaffolded antigens,^41–43^ since they are in the “beyond-rule-of-five” chemical space^44^ and have many protein characteristics.^45^ Computational methods have been developed to design well-structured cyclic peptides that preferentially populate a single conformation,^45–50^ which have led to several applications.^51–53^ However, most cyclic peptides reported thus far have labile structures and adopt multiple conformations in solution. ^54–58^ Although some algorithms excel in creating a diverse ensemble of cyclic peptide conformations,^59–65^ they seldom address the entropic contribution of a flexible polymer, and in practice they often do not provide an ensemble with reliable thermodynamic properties. More recently, enhanced MD methods such as metadynamics and REMD have had success in generating accurate conformational ensembles.^32,66–69^ In this study, we systematically applied Res-REMD on several cyclic peptides, to examine how Res-REMD might improve sampling over other simulation methods such as REMD and conventional MD.

Since this research is the first to incorporate Res-REMD to the GROMACS platform, we started by validating the implementation with a simple system, the Alanine dipeptide. After that, we performed Res-REMD on Trpzip2 to confirm its efficiency over regular REMD,^35^ now using explicit solvent. We also performed multiple additional Res-REMD simulations on Trpzip2 to explore the effect of different reservoir compositions on the final ensemble generation. Finally, we applied Res-REMD on three cyclic peptide systems.

We found that conventional molecular dynamics (MD) simulation was insufficient to sample the ensemble of small cyclic peptides (*<* 10 amino acids), likely due to energy barriers rooted in the ring-like conformational constraint. Enhanced MD methods such as REMD or Res-REMD methods are thus required to sample the complete cyclic peptide ensemble, as previous studies have suggested. ^32,66–69^ For cyclo-(CGPRRARSG) and oxytocin, all REMD and Res-REMD simulations generated similar ensembles for the 300K replica, and these were distinct from the conventional MD ensemble. Moreover, we found that Res-REMD could significantly accelerate equilibrium ensemble generation for cyclo-(CGHHQKLVG) compared to regular REMD. To explain whether or not Res-REMD can improve sampling performance compared to REMD on cyclic peptides, we found that the torsional angle auto-correlation time could be an indicator to determine which cyclic peptide systems would benefit more from Res-REMD.

## 2 Method

### 2.1 Model systems

There are five model systems as shown in Fig. 1: (1) Alanine dipeptide, an ACE-ALA-NME construct (i.e. terminally blocked alanine); (2) Tryptophan zipper 2, or Trpzip2 (PDB ID 1LE1^70^); (3) Oxytocin (PDB ID 2MGO^71^), a disulfide-cyclized 9-mer peptide with sequence CYIQNCPLG; (4) Cyclo-(CGPRRARSG), a backbone-cyclized 9-mer peptide; (5) Cyclo-(CGHHQKLVG), a backbone-cyclized 9-mer peptide.

**Figure 1:**
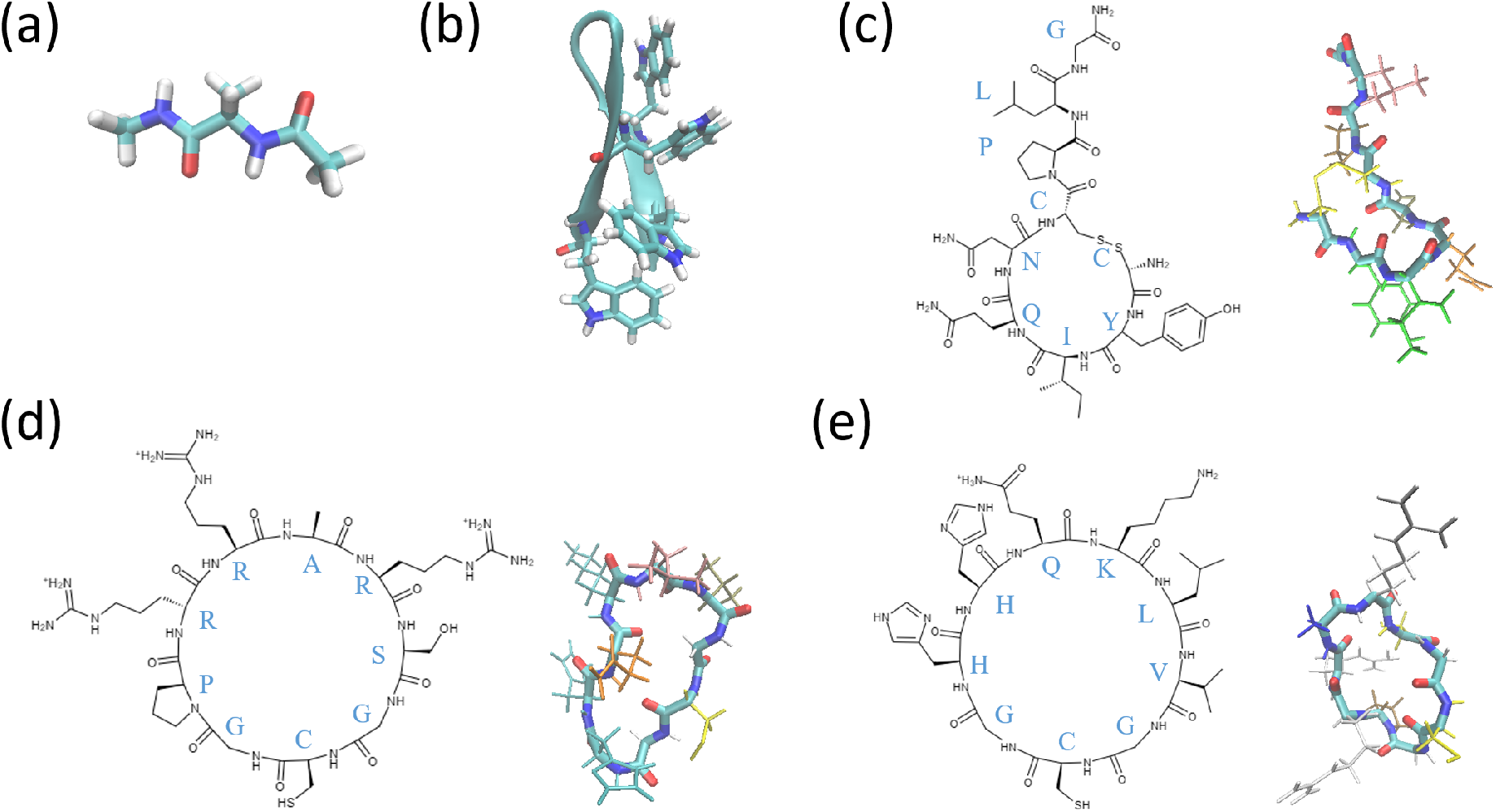
Structural representations of five protein systems under study. For cyclic peptides, the amino acid identities are shown beside their position in the 2D representations. The initial conformation of each cyclic peptides are shown. The backbone color is rendered by atom identities, and each residue sidechain is rendered in a different color. (a) Alanine dipeptide (b) Trpzip2. Tryptophan residues are rendered in licorice to show their close interaction. (c) Oxytocin (d) cyclo-(CGPRRARSG), cyclized furin cleavage site of SARS-CoV-2 spike protein (e) cyclo-(CGHHQKLVG), cyclized segment of A*β* protein.

### 2.2 Protein system choice

Alanine dipeptide is a simple molecule, readily used for testing simulation techniques and free energy methods. Trpzip2 is an artificially designed 12-amino-acid protein with a high propensity to form a beta-hairpin in solution at room temperature. The temperature-dependent folding stability of Trpzip2 provides an excellent model for testing sampling algorithms. Oxytocin is a sidechain-cyclized peptide hormone produced in the brain, and is also available in pharmaceutical form. Oxytocin has also been well-characterized by NMR experiments,^72^ which allows us to compare the computationally generated ensemble against the NMR measurements.

Cyclo-(CGHHQKLVG) is a glycine-flanked and cyclized epitope for A*β* protein,^43^ where the sequence HHQKLV was predicted to be misfolding-prone ^73^ by a collective coordinate biasing method.^74^ Cyclo-(CGHHQKLVG) in conjugation with carrier protein (e.g. KLH or BSA) is an immunogen candidate, and similar cyclic peptides (cyclo-(CGHHQKG)) have been used as immunogens to raise conformationally selective antibodies to A*β* oligomers.^42^ Similar scaffolds of other epitopes have also been used to successfully raise oligomer-selective antibodies for Alzheimer’s disease. ^41^ The scaffolding method by adding a variable number of flanking glycines and a cysteine (the “glycindel” method) follows the rationale described previously.^43^ In brief, flanking glycines add more flexibility to the scaffold, and induce antibodies that recognize more diverse conformations of the epitope; The flanking cysteine is required to conjugate the cyclic peptide to the carrier protein through a disulfide bond.

Cyclo-(CGPRRARSG) is a scaffolded epitope for SARS-CoV-2 spike protein, where the sequence PRRARS is at the S1/S2 domain cleavage site. An antibody that is raised by this cyclic peptide antigen could potentially inhibit the cleavage, and have a neutralization effect.^75^ An antibody recently raised by the KLH-conjugated SARS-CoV-2 peptide (^672^ASYQTQTNSPRRARSVASQS^691^) was observed to efficiently block S1/S2 cleavage and binding to ACE2.^76^ Note that the proline in our epitope was identified to have key mutation in Alpha variant (P681H) and Delta variant (P681R), which significantly increased viral replication and transmission,^77,78^ largely due to increased cleavage of S1/S2 site by the furin enzyme.^76^

### 2.3 General simulation setup

#### 2.3.1 Input structure model

The Alanine dipeptide structure contains three residues ACE-ALA-NME, and was taken from previous studies.^36,79^ The native structures for Trpzip2 and oxytocin were taken from the first NMR models for each respective structure, 1LE1^70^ and 2MGO.^71^ The unfolded structure of Trpzip2 was taken from a random snapshot of a conventional MD simulation at 416K that fulfills RMSD from the backbone of an energy minimized native structure *>*0.29nm. Visual inspection had further confirmed the structure was extended. For cyclo-(CGHHQKLVG) and cyclo-(CGPRRARSG), the structures were prepared using a method described previously.^43^ The structure of the HHQKLV motif was taken from PDB ID 2M4J, ^80^ and the structure of the PRRARS motif was taken from the SARS-CoV-2 spike protein structure released by Deepmind on August 4 2020.^81^ The histidine protonation states in cyclo-(CGHHQKLVG) were all HSE (protonated on epsilon nitrogen, NE2), which were assigned by the optimal hydrogen bonding pattern calculated by the pdb2gmx module. For Alanine dipeptide, no modification was performed on the termini, since the construct is already terminally blocked by ACE and NME residues. For Trpzip2, both termini were charged (NH ^+^ and COO^−^). For oxytocin, the N-terminus was charged (NH ^+^), and the C-terminus was amidated (NH2), consistent with the modifications present in experiments.^71,82^

#### 2.3.2 Minimization and Equilibration

Simulations were run using GROMACS-4.6.7 with a custom Res-REMD implementation and the CHARMM36m force field.^83^ A dodecahedron unit cell with box boundary 1.2 nm distance away from the closest atom on the protein was used for simulations, wherein each model system was solvated with explicit TIP3P water,^84^ and Na^+^ and Cl^−^ ions were added to neutralize the system charge and maintain an aqueous salt concentration of 100 mM (150mM for cyclic peptides). System energy was then minimized through steepest descent until a maximum force *<* 100 kJ/mol/nm, followed by 100 ps NVT thermostat through the velocity-rescale method, with 1000 kJ/mol/nm positional restraints on the heavy atoms. Protein and solvent thermostats were performed separately, and have a coupling time of 0.1 ps. A time step of 2 fs was used in all simulations. NVT equilibration was followed by a 300 ps NPT thermostat using Parrinello-Rahman and velocity-rescale method with 1000 kJ/mol/nm positional restraints on heavy atoms. The pressure coupling is isotropic with a coupling time of 2 ps, and compressibility of 4.5 *×* 10^*−*5^ bar^*−*1^. Electrostatics was calculated by the PME method with order 4, and Fourier spacing 0.16. The electrostatics cutoff and van der Waals cutoff were both 1.0 nm. LINCS constraints method of order 4 was applied on heavy atom-hydrogen covalent bonds, with iteration set to 1. The temperature and pressure were maintained at 300 Kelvin and 1.0 bar, respectively.

#### 2.3.3 REMD simulation details

Testing the efficiency and accuracy of Res-REMD simulations requires a reference simulation, because the experimental data cannot yield the “correct” answer using a *given* force field and solvent model. Regular REMD simulation was thus performed to compare with Res-REMD. The temperature interval between REMD replicas was calculated from the server https://github.com/dspoel/remd-temperature-generator,^85^ to achieve a target average exchange rate of 0.2/swap attempt between all neighboring replicas. Before MD production runs, short 100 ps NVT and 300 ps NPT equilibration runs were performed at each target temperature to equilibrate each replica. Exchanges between replicas were attempted every 1 ps for all simulations. The exchange attempt alternated between even and odd replica pairs (i.e. the highest and the lowest replica performed the exchange attempt every 2 ps). Parameters for each REMD simulation are listed in Table 1. Snapshots from simulations were taken every 10 ps; These structures are used to obtain a fraction of the ensemble that is either in a given region of Ramachandran space (Alanine dipeptide), a folded fraction (Trpzip2), or the torsional entropy (Cyclic peptides).

**Table 1:**
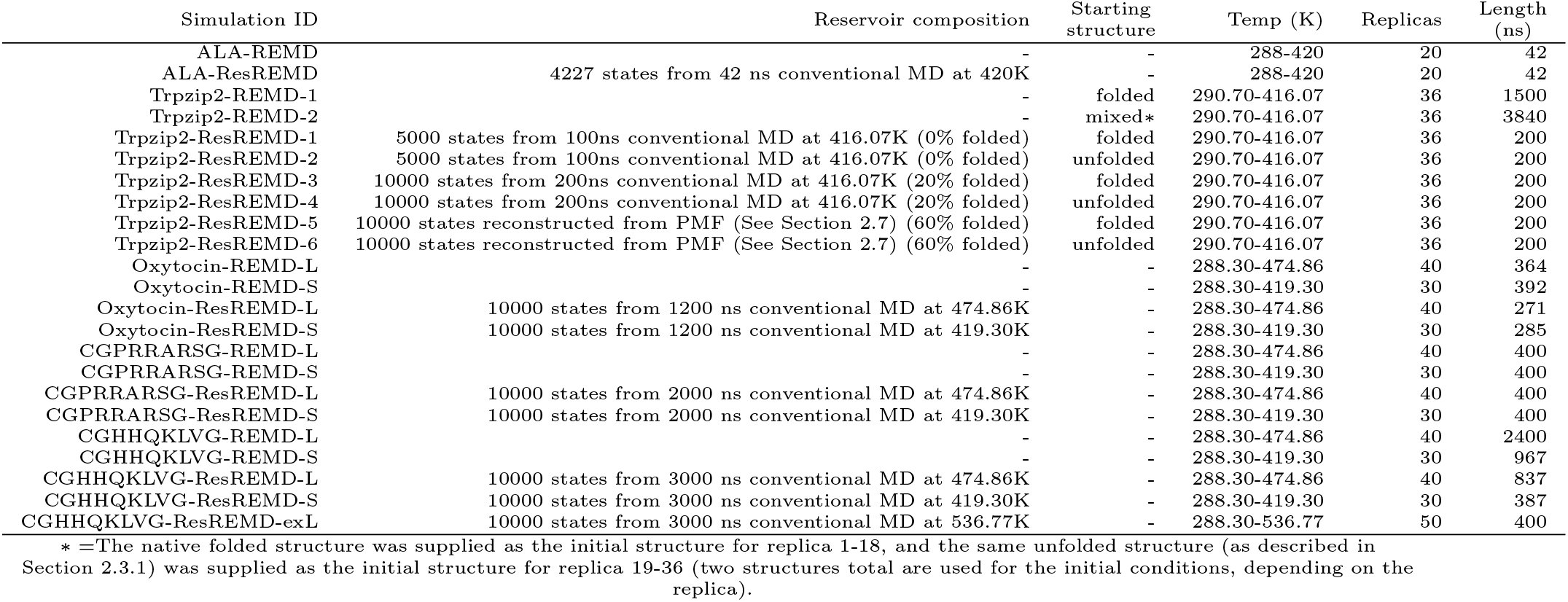
Res-REMD and REMD simulations in this study

#### 2.3.4 Res-REMD simulation details and Reservoir generation

Res-REMD simulations also used the same procedure as REMD simulations, except that in addition to the exchanges between replicas, exchanges between the highest temperature replica and the reservoir were attempted every 2 ps. The reservoir of Trpzip-ResREMD-5 and Trpzip2-ResREMD-6 were generated based on Potential of Mean Force (PMF) along the unfolding coordinate, RMSD against the native structure, using umbrella sampling (Section 2.7). For all other systems except for Trpzip2-ResREMD-5 and Trpzip2-ResREMD-6, reservoir generation was performed at a high temperature using conventional MD (see Table 1). All Res-REMD simulations in this study are Boltzmann-weighted Res-REMD, in that reservoir is Boltzmann sampled. The number of states in the reservoir for each simulation are all well above the recommended range of 1000-5000 conformations^86^ for a Boltzmann-weighted reservoir.

### 2.4 Reservoir-REMD implementation in GROMACS

The GROMACS 4.6.7 source code was modified to implement Res-REMD. We made use of checkpoint files, which are saved regularly in GROMACS for the purpose of continuing the simulation, to build the reservoir. The checkpoint file contains all the information necessary to continue a simulation, which can be extracted from the trajectory files (i.e. .trr file). We edited the GROMACS 4.6.7 source code to implement the following processes:

1. Before each replica exchange attempt, the highest-temperature replica (i.e. the reservoir) reads a random checkpoint file. This was followed by a single 2 fs MD step, and the state is halted while the other replicas continue to perform MD integration until the next replica exchange was attempted. While the state in the reservoir replica was halted, the message passing interface (MPI) communication with other replicas was kept to synchronize global communications (e.g. checkpoint saving, programmed termination). Potential energy was calculated within the single MD step as the exchange criterion.
2. Since processing the reservoir replica is a simple task, the MPI architecture was modified so that the reservoir replica only took one CPU core. At this stage of our development, Res-REMD can only run on CPU, and GPU support was not added.
3. A new mdrun flag, -reservoir *<*\*cpt files*>*, was added. Once this flag is issued, the highest temperature replica becomes the reservoir, and the given checkpoint files are read in as the reservoir configurations.

Many technical modifications were made, but only core modifications are as described above. The full code and instructions are available on Github: https://github.com/PlotkinLab/Reservoir-REMD.

### 2.5 Convergence Analysis

The convergence of simulations was determined by three different methods respectively applied to each class of model system: Alanine dipeptide, Trpzip2, and cyclic peptides. The definition of these metrics are described below.

#### 2.5.1 Alanine dipeptide: binning of Ramachandran space

Using the same definition as Henriksen et al.,^36^ the Ramachandran space of Alanine dipeptide was binned into 6 macrostates (Fig. 2a). Convergence was observed as the saturation of the frequency of occurrence for each macrostate versus simulation time.

**Figure 2:**
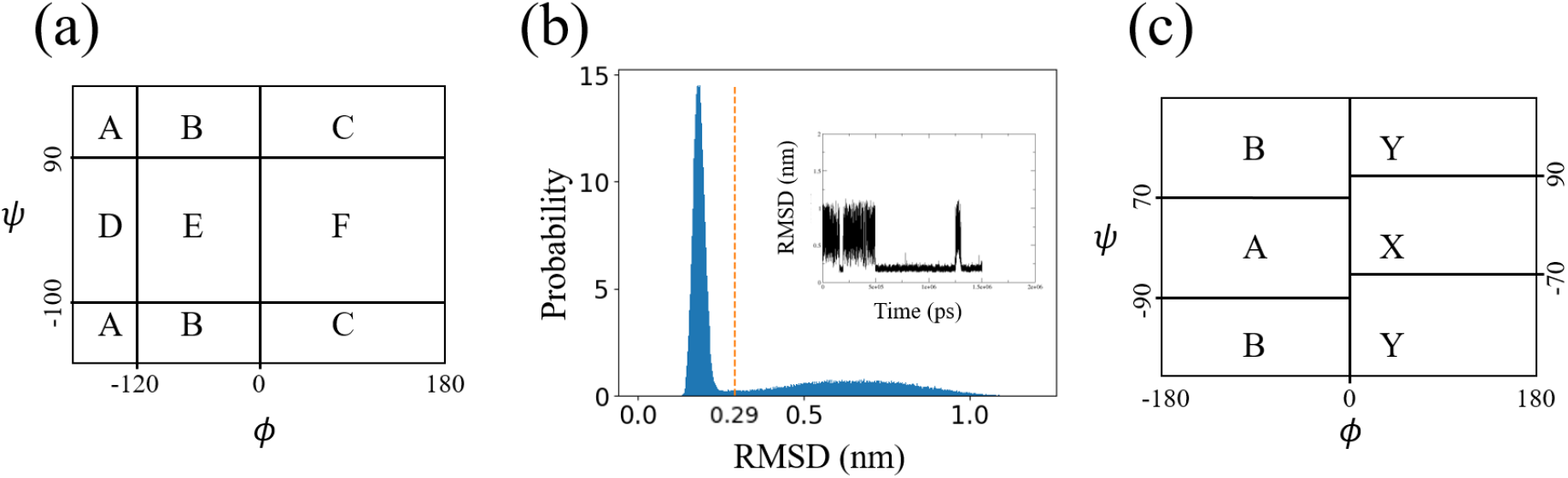
Definitions of the coarse-grained states applied to each class of model system. (a) For Alanine dipeptide, 6 coarse-grained states A-F are defined by binning the space of Ramachandran angles. (b) For Trpzip2, coarse-grained folded and unfolded states are defined by an RMSD cut-off at 0.29nm that separates the states. The location of the RMSD cut-off was determined by the local minimum of the RMSD histogram after 1500 ns of conventional MD simulation at 416.07K. The inset shows the folding/unfolding trajectory projected onto RMSD. (c) For cyclic peptides, macrostates A, B, X, and Y states are defined by binning the Ramachandran space for each amino acids.

#### 2.5.2 Trpzip2: folded fraction

As mentioned above, RMSD was calculated against backbone heavy atoms of all 12 residues in an energy minimized native structure. The RMSD cutoff for separating the folded and unfolded macrostate is 0.29 nm, which corresponds to the local minimum of the RMSD frequency histogram (Fig. 2b). Convergence was observed as a saturation of the folded fraction verses simulation time.

#### 2.5.3 Cyclic peptides: torsional entropy

Similar to the Ramachandran space binning for Alanine dipeptide, the torsional angle of each amino acid in CYIQNC, PRRARS, and HHQKLV (respectively for model system 3-5) was assigned a coarse-grained macrostate from A, B, X or Y in the Ramachandran space (Fig. 2c). This definition of ABXY torsional bins was first proposed by Hosseinzadeh et al.^50,87^ A and B represent the backbone Ramachandran space of alpha-helix and beta-sheet, respectively; X and Y are the mirror of A and B (i.e. the alpha-helix and beta-sheet for D-amino acids). For example, the coarse-grained state of HHQKLV in cyclo-(CGHHQKLVG) can be represented by a 6-letter string for the torsional angle bins of H, H, Q, K, L, and V, respectively (e.g. AAAAAA, ABAABX, etc…). The total number of states defined by ABXY torsional bin is 4^6^ = 4096 (4 torsional bins to the power of 6 amino acids). The torsional entropy was used to determine the convergence of cyclic peptide simulations; Torsional entropy was defined as 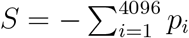, where i runs over all possible torsional macrostates. When plotted verses simulation time, the flattening of entropy in a simulation trajectory suggests that the structural diversity of the epitope is no longer changing over time. The entropy measure here has been called the Shannon diversity in other studies ^88^ measuring the diversity of component states.

### 2.6 Ensemble similarity analysis for cyclic peptides

#### 2.6.1 Torsional Similarity

In addition to defining the torsional entropy, the set of torsional state probabilities {*p*_*i*_} can also be compared between ensembles to obtain an ensemble similarity. The torsional similarity is the sum of the square root of the dot product of the torsional state probabilities 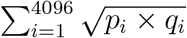, where *i* indicates a given torsional state defined by all angles in the epitope. This method is similar to finding the cluster population correlation from combined clustering.^86^ The difference here is that our clusters were pre-defined to be the torsional states, instead of being assigned from structural clustering.

#### 2.6.2 Jensen-Shannon Divergence

Since torsional similarity involves segmentation of the Ramachandran space, structures close to the boundary could be assigned to different torsional states despite being structurally similar. To resolve this issue, we also calculated the Jensen-Shannon Divergence (JSD),^89–91^ a similarity measure that does not require segmentation of the Ramachandran space. JSD was implemented here using the ENCORE software.^92^ JSD is a symmetrized and smoothed version of the Kullback–Leibler divergence,^93,94^ *D*_*KL*_, which is a difference measure between two distributions. JSD ranges from 0 to log(2), and a larger value indicates less similarity. The JSD calculation depends on the dimension of the reduced space into which the pairwise RMSD matrix is projected. Stochastic proximity embedding^92^ was performed on the RMSD matrix to reduce the dimension from (*N × N*) to (*N × D*), where *D* is the desired lower dimension. A reduced dimensional space of *D* = 5 was used for measuring the ensemble similarity. The method of utilizing JSD to measure the ensemble similarity was also employed in our previous work. ^43^

Structural ensembles were extracted from the 300K replica with 500 ps sampling interval for CGPRRARSG and CGHHQKLVG, and 150 ps sampling interval for oxytocin. Ensembles of conventional MD were extracted every 1 ns. The time interval was confirmed to be larger than the RMSD auto-correlation time in each simulation trajectories. Only heavy atoms in CYIQNC, PRRARS, and HHQKLV (model system 3-5 respectively) were included for ensemble comparison.

### 2.7 Reservoir construction using potential of mean force (PMF) for Trpzip2

The reservoir for Trpzip2 was generated at 416.07 K, well-above the reservoir temperature of 400 K in a previous Trpzip2 study.^28^ To acquire a reservoir with Boltzmann-weighted component states, we used a PMF-weighted method involving Umbrella Sampling (US). The method is composed of two steps, detailed below. Note that the method used here is computationally inefficient for the reservoir generation problem, since one may have used US to directly build Potential of Mean Force (PMF) at the target temperature (e.g. 300K). In general, US may not be computationally efficient to build reservoir in most real applications.

1. **PMF construction:** The PMF along RMSD was constructed using the MBAR^95^ estimator on the US trajectories. US was performed serially — i.e. the final coordinate of a given US window serves as the initial coordinate of the next US window. Thus, each successive window benefits at least partially from a longer equilibration time.^96^ The simulation took 5 ns for each of the 55 windows generated during serial window construction. Subsequently, each window was extended for another 35 ns (Fig. 3a). The resulting PMF shows two local minima with a free energy difference of about 2 kcal/mol (Fig. 3b). The energy barrier located at 0.29 nm recapitulates the RMSD cutoff defined previously using conventional MD. The folded state (RMSD *<* 0.29 nm) consists of approximately 60% of the reservoir.
2. **PMF-weighted reservoir construction:** The reservoir was constructed based on the PMF. All structures saved during the last 50% of US trajectories were classified into RMSD bins (e.g. RMSD equals 0.2-0.25 nm). The reservoir was then constructed from randomly drawn states at each RMSD bin, wherein the number of structures drawn from each RMSD bin *j* is proportional to the corresponding Boltzmann factor 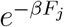 for that bin; *F*_*j*_ is the PMF value. As expected, the reservoir RMSD distribution generated by this procedure was confirmed to be the same as the Boltzmann distribution calculated from inverting the PMF appearing in the exponential (Fig. 3c).

**Figure 3:**
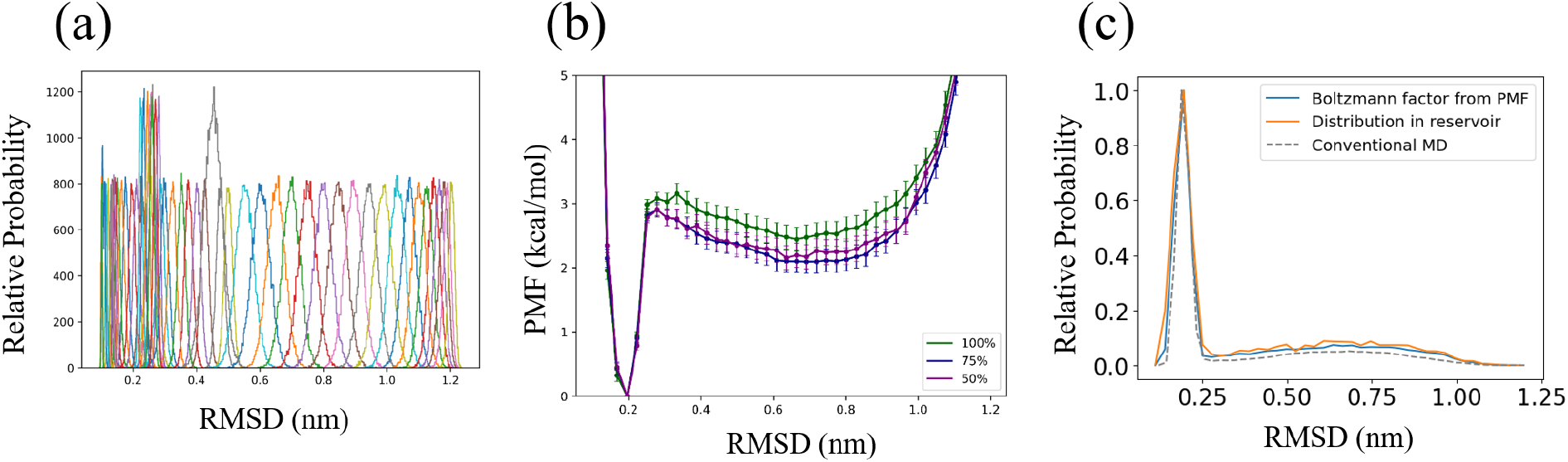
The procedure and analysis of PMF construction for Trpzip2. (a) The RMSD histograms for all US windows. (b) The PMF of Trpzip2 along RMSD at the reservoir temperature of 416.07K. The convergence is addressed by the similar PMFs using the last 75% of the US trajectories and the last 50% of the US trajectories. (c) The reservoir RMSD distribution obtained from the procedure in Section 2.7 agrees with the distribution calculated by directly inverting the PMF appearing in the exponential, using the last 50% US trajectories. For comparison, the distribution of RMSD extracted from direct MD simulations (Fig. 2b) is shown as the dashed curve.

### 2.8 Other analyses

#### 2.8.1 t-Distributed Stochastic Neighbor Embedding (t-SNE)

The dimensional reduction using t-SNE^97^ analysis was used to visualize the reservoir phase space projected to 2D (see e.g. Fig. 7b). The phase space was represented by a pairwise RMSD matrix, in which the structures were evenly drawn from the simulation trajectory. Then, t-SNE reduced the *N ×N* matrix to *N ×*2. t-SNE was implemented by the scikit-learn module in Python.

#### 2.8.2 time-lagged Independent Component Analysis (tICA)

tICA analysis can find independent slow reaction coordinates.^98,99^ With a lag time of 10 ns, tICA was used to find a linear combination of cosine of torsional angles that has the slowest reaction coordinate as represented by the normalized first eigenvector. tICA was implemented by pyemma module in Python.

## 3 Results and Discussion

### 3.1 Validitation using Alanine dipeptide

The free energy profiles of the 300K replicas were extracted from the ALA-REMD and ALA-ResREMD simulations by the histogram method^100^ (Fig. 4a,b). The histogram method estimates the probability in Ramachandran space by Kernal Density Estimation (KDE), and the relative free energy was calculated from the relation *F* (*φ, ψ*) = *−k*_*B*_ ln (*p* (*φ, ψ*) */p*_*Max*_). Both simulations have the same location of global free energy minimum, in the upper-left quadrant of the Ramachandran space (in macrostate B of the coarse-grained binning scheme described in the Section 2.5.1.) Moreover, the frequency of occurrence of macrostates A-F for the 300K replica for both ALA-REMD and ALA-ResREMD converged to the same value within 42 ns (Fig. 4c). The convergence of the reservoir generation process was also confirmed by the flattening of the occurrence frequency of each state as the simulation progresses. (Fig. 4d). Note that the probabilities of each coarse grained state are more uniform at the higher temperature of the reservoir (c.f. Fig.s 4c,d). The convergence of the free energy surface using either REMD or Res-REMD supports the validity of the GROMACS implementation of Res-REMD introduced here.

**Figure 4:**
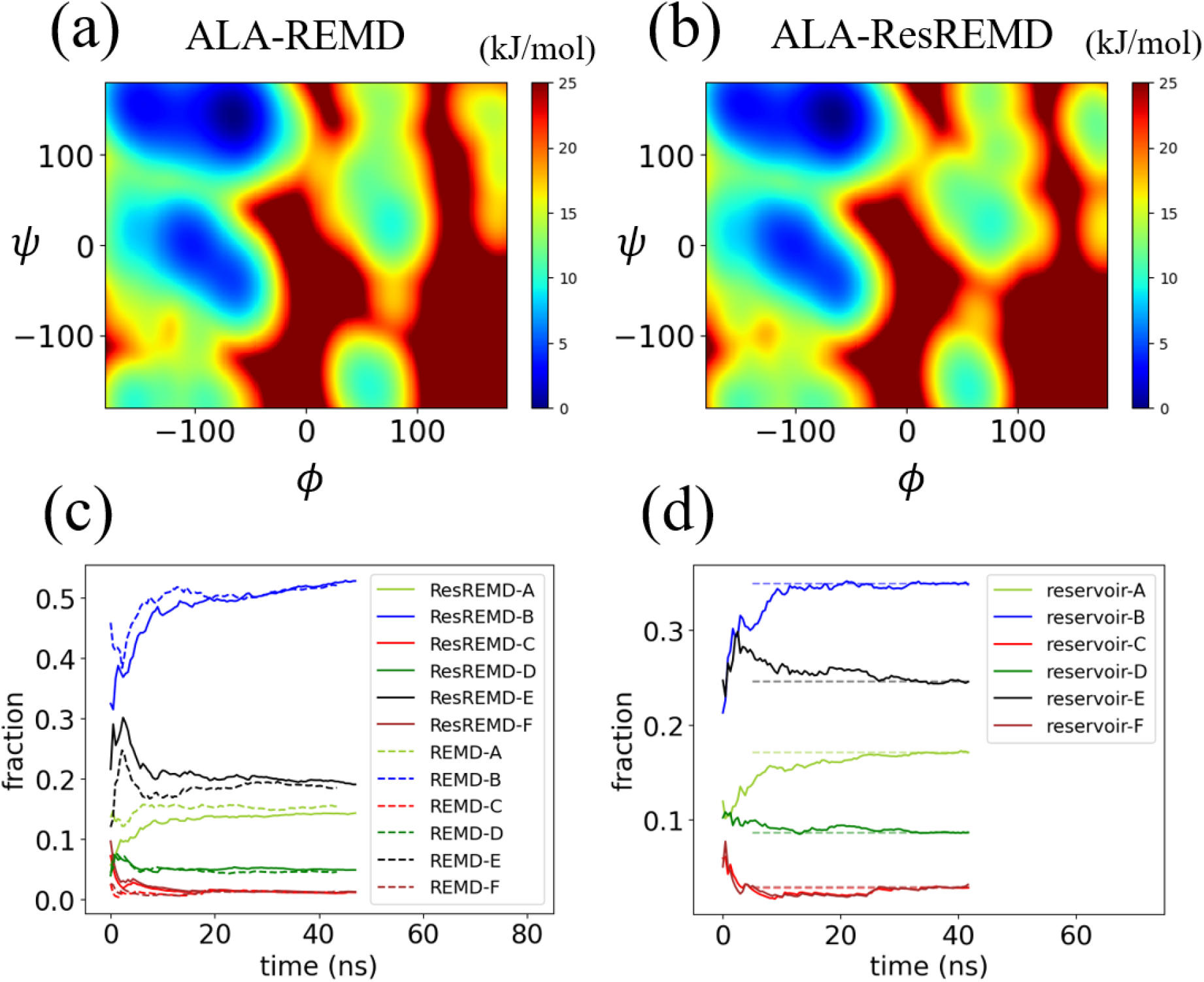
(a) The free energy profile of Alanine dipeptide using (a) REMD and (b) Res-REMD. (c) The frequencies of occurrence for the six coarse-grained states (A-F as defined in Fig. 2a), for the 300K replica of both ALA-REMD and ALA-ResREMD. (d) The fraction of total states contained in each macrostate A-F, in the T=420K reservoir, for ALA-ResREMD. Time here corresponds to the samples taken from the reservior as the simulation progresses. The asymptotic values determined by the composition of the reservoir are also shown as dashed lines.

### 3.2 Application of Res-REMD to Trpzip2 in explicit solvent

It has been shown by Kasavajhala et al. ^86^ that correct ensemble generation in Res-REMD is contingent upon an accurate Boltzmann-weighted reservoir. To generate an correct reservoir for Trpzip2, we first attempted conventional MD. However, there were only 5 folding/unfolding transitions observed within the total simulation time of 1500 ns (Fig. 2b inset). This is far from the 100 folding/unfolding transitions suggested for sufficient reservoir convergence.^86^ To generate a correct reservoir, a PMF-weighted method was thus used (See Section 2.7).

To test the efficiency and accuracy of Res-REMD simulations against reference data, we performed conventional REMD simulations (See Section 2.3.3) starting from different initial conformations (Table 1). All REMD and Res-REMD simulations spanning the same temperature range have the same replica temperatures, which enabled direct comparison in each replica temperature (see Fig. 5 and description below).

**Figure 5:**
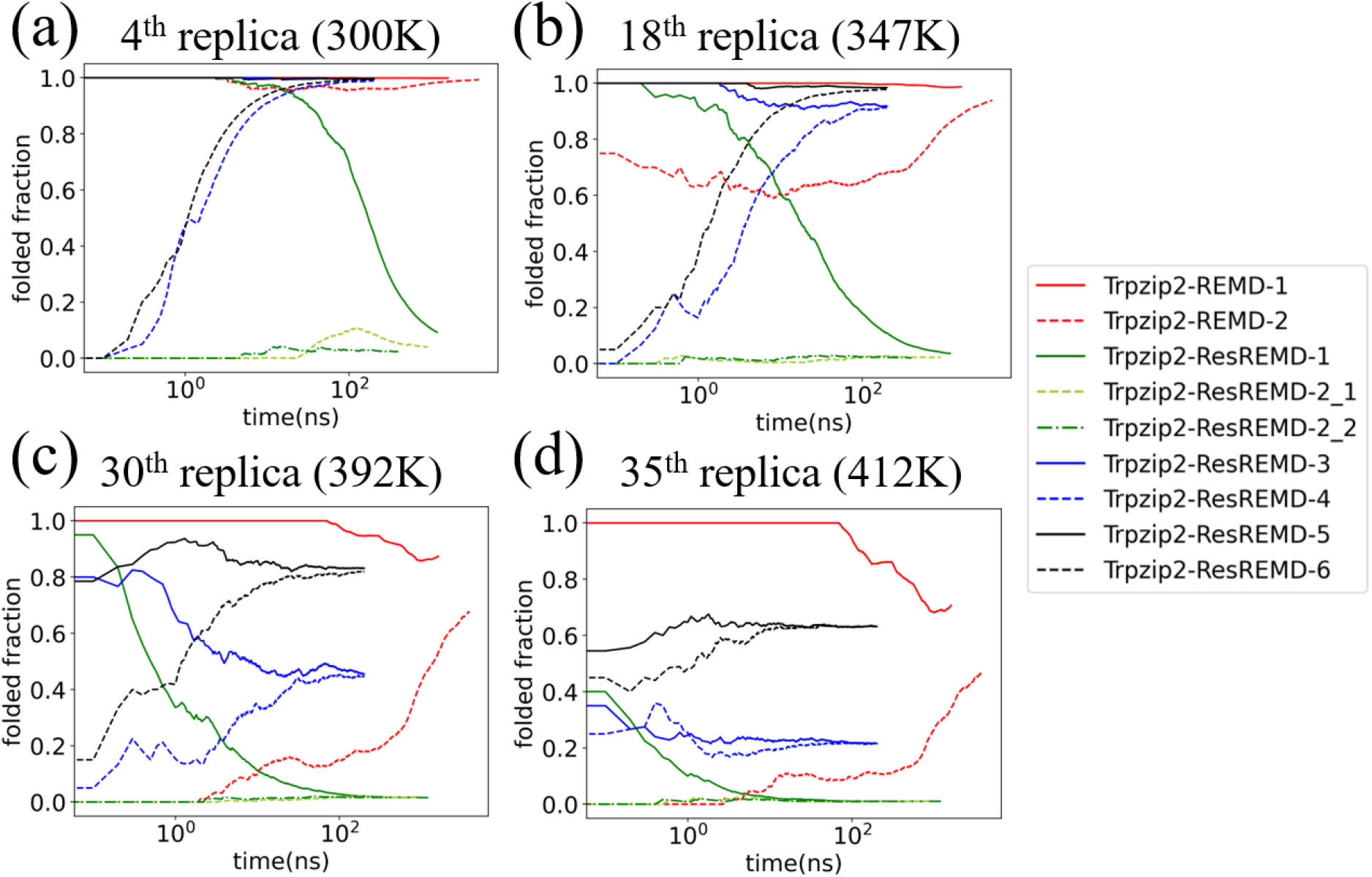
The folded fraction of all the Trpzip2 simulations in four different replicas, at 4 different temperatures. (a) 4^th^ replica (300K) (b) 18^th^ replica (346.71K) (c) 30^th^ replica (391.70K) (d) 35^th^ replica (411.91K). The folded fraction is obtained as a running average with an interval of 100 ps; The initial value of the folded fraction is determined from an ensemble collected in the first 100 ps (i.e. the first 10 conformations).

#### 3.2.1 Res-REMD is robust to different initial conditions, but not the change of the reservoir composition

To understand how the composition of the reservoir affects Res-REMD simulations, three reservoirs were extracted for Res-REMD simulations of Trpzip2. The first reservoir is composed of 5000 states from the first 100 ns of a conventional MD simulation, and has 0% folded fraction. The second reservoir is composed of 10000 states from the first 200 ns of a conventional MD simulation, and has 20% folded fraction. The third reservoir is composed of 10000 states from a PMF-weighted reservoir construction method (Section 2.7), and has 60% folded fraction. The first reservoir was coupled to simulations Trpzip2-ResREMD-1 and Trpzip2-ResREMD-2 (two repeats of Trpzip2-ResREMD-2 were labeled with suffix 1 and 2), the second reservoir was coupled to simulations Trpzip2-ResREMD-3 and Trpzip2-ResREMD-4, and the third reservoir was coupled to simulations Trpzip2-ResREMD-5 and Trpzip2-ResREMD-6 (Table 1). Initial conditions of simulations are also given in Table 1.

Even with different initial conformations, Res-REMD simulations that were coupled to the same reservoir converged to the same folded fractions (Fig. 5). However, Res-REMD simulations using different reservoirs converged to different folded fractions. Therefore, Res-REMD is robust to different initial conditions, but not changes of reservoir composition. Thus, a correctly Boltzmann-weighted reservoir is essential for generating correct Res-REMD simulations. Moreover, incorrect reservoirs result in an incorrect equilibrated ensemble, as described further below in Section 3.2.3.

#### 3.2.2 Res-REMD simulations show higher efficiency than REMD simulations

Two REMD simulations with different initial conditions were performed, which were observed to exhibit much slower convergence than Res-REMD simulations. The slow convergence of REMD simulations can be seen particularly in Fig. 5c,d, where the folded fraction in Trpzip2-REMD-1 and Trpzip2-REMD-2 slowly approaches that of the already converged Res-REMD simulations having a correctly Boltzmann-weighted reservoir (Trpzip2-ResREMD-5 and Trpzip2-ResREMD-6). Trpzip2-REMD-1 barely converges to the same folded fraction as the correct Res-REMD simulations (Fig. 5), while Trpzip2-REMD-2 slowly approaches the correct folded fraction of the Res-REMD simulations, but without achieving convergence within 3.84 *µ*s. This was more than twice the simulation time of Trpzip2-REMD-1 (note the log scale of Fig. 5). The faster apparent convergence of Trpzip2-REMD-1 than Trpzip2-REMD-2 is likely due to its closer initial condition to the equilibrated ensemble.

Note that, at the second-highest temperature replica #35 at T=412K (Fig. 5d), both REMD simulations were expected to converge to about 60%, which is the folded fraction of the correct reservoir. Consequently, Trpzip2-REMD-1, with folded initial structures in all replicas, is somewhat closer to the equilibrium ensemble than Trpzip2-REMD-2, where half of the replicas begin with unfolded structures. The higher efficiency of Res-REMD than REMD agrees with a previous Trpzip2 study using implicit solvent.^28^

#### 3.2.3 Incorrect convergence of Res-REMD when the reservoir is far from Boltzman-weighted distribution

By varying the reservoir composition, we observed that an incorrect reservoir far from the Boltzmann-weighted distribution could induce an incorrect equilibrated ensemble, which may also exhibit slow convergence. For example, coupling to a 0% folded fraction reservoir of Trpzip2-ResREMD-1 induces a slower convergence compared to Trpzip2-ResREMD-{3,4,5,6}, which is most dramatic in Fig. 5b. Most importantly, the convergence of Trpzip2-ResREMD-{1,2,3,4} approaches the incorrect folded fraction in Figure 5. The reservoir in the Res-REMD simulations may be thought of as a boundary condition on the sampling trajectories. The boundary condition determines the converged, equilibrated ensemble at 300K, regardless of the initial condition. An incorrect boundary condition (e.g. a non-Boltzmann weighted reservoir) leads to the generation of an incorrect converged ensemble.

We also found, perhaps not surprisingly, that a Res-REMD simulation can generate structures at lower temperatures not present in the reservoir. Specifically, with no folded structure in the reservoir (Table 1), Trpzip2-ResREMD-2 1 still shows folding events, resulting in a transient increase in the folded fraction from 0 to 0.1 at around 100ns in the low-temperature replica (light green dotted line in Fig. 5a). However, Trpzip2-ResREMD-2 2, a repeat of Trpzip2-ResREMD-2 1, shows low folded fraction throughout the simulation. Thus, the spike of folded fraction in Trpzip2-ResREMD-2 1 was caused by a stochastic folding event. A detailed analysis for Trpzip2-ResREMD-2 1 (Fig. S1) shows that a folding event occurred at around 22ns, and the protein stayed folded during 22-130ns, which contributes to the spike in the folded fraction.

#### 3.2.4 Comparison to the experiment

The experimentally measured melting temperature for Trpzip2 is 345K,^70^ where the folded fraction is 50%. On the other hand, the folded fraction in the PMF-weighted reservoir (at T=416K) only achieved 60%, which suggests that the Trpzip2 melting temperature is larger than this reservoir temperature. The reason for this discrepancy may be two-fold. Firstly, Trpzip2 in this study has charged N- and C-termini (NH ^+^–SWTWENGKWTWK– COO^−^), but the experimental construct differs, in that it has a neutral amidated C-terminus (NH ^+^–SWTWENGKWTWK–CONH).^70^ The negatively charged Trpzip2 C-terminus in simulations may lead to an increased melting temperature, due to attractive forces with the positively charged N-terminus. Secondly, the force field will have an effect on the folding stability and melting temperature of Trpzip2. Although past simulations using Amber ff99 force field with modified backbone parameters ^101^ and generalized Born implicit solvent^102^ had successfully demonstrated that the melting temperature of Trpzip2 is close to 345K,^28^ the Trpzip2 simulation in this study using the CHARMM36m force field with TIP3P water has not been performed previously, and may not be able to reproduce the correct melting temperature even for the identical system in experiments.

### 3.3 Cyclic peptides

#### 3.3.1 Conventional MD did not generate a converged ensemble at 300 K

Conventional MD simulations at 300K were attempted first to generate the ensembles for the cyclic peptides, but none of them achieved convergence (Fig. 6a-c black lines). A converged ensemble for cyclic peptide was defined here as the flattening of the torsional entropy curve (See Section 2.5.3). The ruggedness of torsional entropy curve for conventional MD at 300K suggests that the cyclic peptides are susceptible to kinetic trapping. On the other hand, conventional MD at higher temperatures (i.e. reservoir temperature at 419K, 475K, and 537K) can generate converged ensembles (Fig. 6d-f). This temperature-dependent convergence property suggests that replica-exchange-based (including both REMD and Res-REMD) simulations can help the 300K replica overcome the energy barriers by exchanging its state with those of a higher temperature replica.^23,25,28^ For this reason, REMD and Res-REMD simulations were performed to accelerate the ensemble generation.

**Figure 6:**
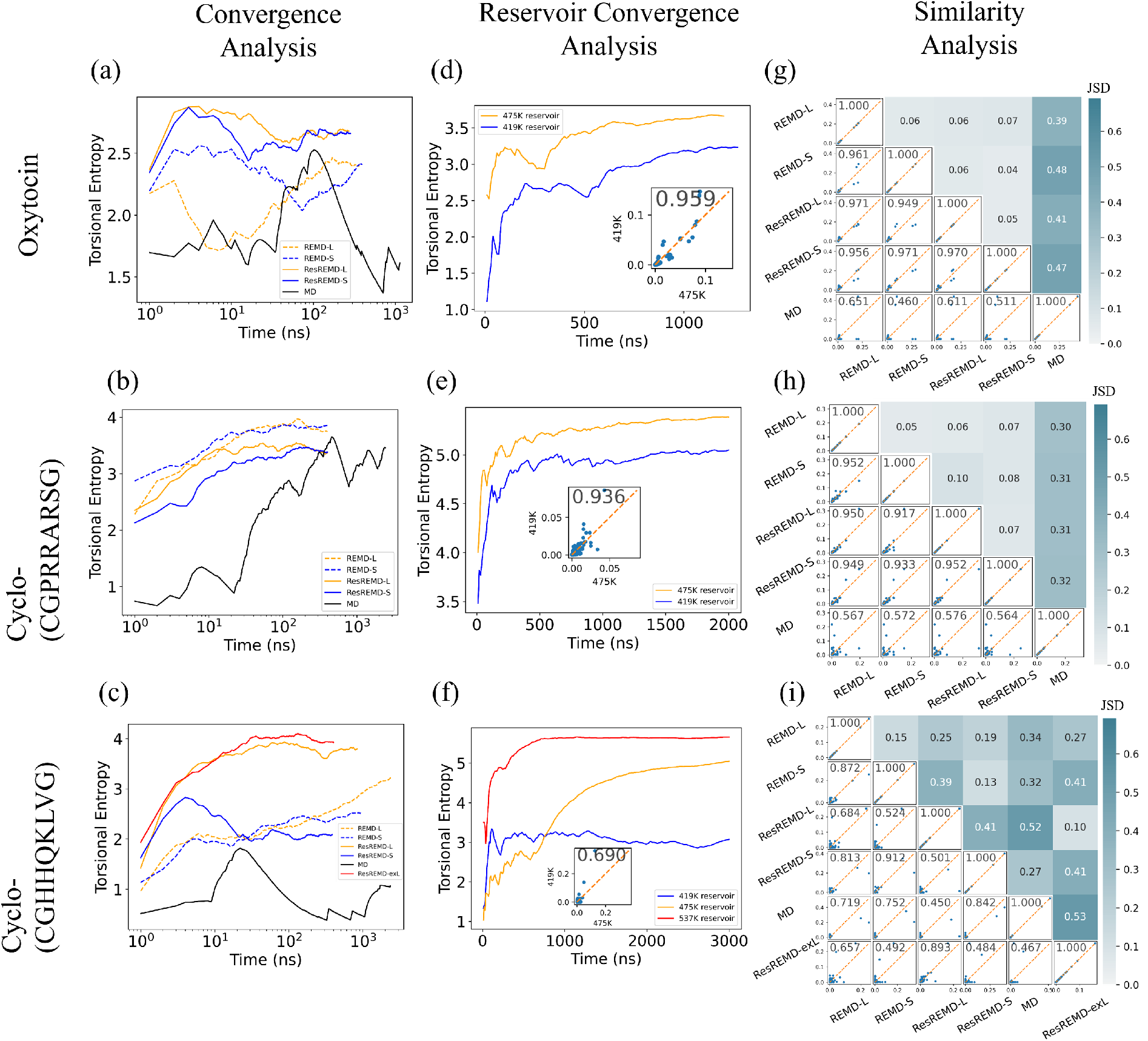
Several analyses on the three cyclic peptides. (a-c) Convergence of the torsional entropy was investigated for the simulated ensembles generated at 300K, for both replica-exchange-based simulations and conventional MD. Cyclo-(CGHHQKLVG) has an additional Res-REMD with a reservoir at 537K. (d-f) These plots show the convergence of torsional entropy during the reservoir generation processes using conventional MD at 475K and 419K. Cyclo-(CGHHQKLVG) has an additional reservoir at 537K. The inset in each panel shows the torsional state probability comparison between the two reservoirs generated at 475K and 419K. (g-i) Two similarity measures were deployed to each pair of ensembles. In the triangle of panels on or below the diagonal, the torsional similarity (Section 2.6.1), is shown at the top left corner of each panel, and in the upper triangle, JSDs were calculated (Section 2.6.2). The blue dots in each scatter plot in the lower triangle represent the fractional population of each of the torsional states; They line up on the x=y line when comparing two identical ensembles (diagonal entries), and there are 4^6^ = 4096 points in each scatter plot, corresponding to each torsional state of the 6 residue epitope.

#### 3.3.2 Res-REMD and REMD produce similar ensembles at similar time scales, and accelerate ensemble generation for oxytocin and cyclo-(CGPRRARSG) over conventional MD

For each cyclic peptide, we performed both REMD and Res-REMD simulations with two different temperature ranges (Table. 1). The simulations with a suffix “-S” or “-L” in Table 1 have an upper bound temperature at 419K or 475K, respectively. For cyclo-(CGHHQKLVG), we performed an additional Res-REMD simulation, suffixed “-exL”, with a higher upper bound temperature at 537K. In sum, a total of 6 REMD simulations (i.e. 3 cyclic peptides *×* 2 temperature ranges) and 7 Res-REMD simulations (i.e. 3 cyclic peptides *×* 2 temperature ranges + 1 CGHHQKLVG-ResREMD-exL) were performed for cyclic peptides.

All the Res-REMD simulations showed smoothing and flattening of the torsional entropy curves (Fig. 6a-c), thus showing evidence of convergence. Care must be taken to ensure this convergence is to the equilibrium ensemble, avoiding incorrect convergence due to coupling to a non-equilibrium reservoir, as dealt with above in Section 3.2.3, and discussed in detail below in Section 3.3.4.

On the other hand, the REMD simulations only partially converged; Oxytocin-REMD-L, CGPRRARSG-REMD-{S,L}, and CGHHQKLVG-REMD-S show more obvious entropy flattening than the other non-converged REMD simulations such as Oxytocin-REMD-S and CGHHQKLVG-REMD-{S,L} (Fig. 6a-c). Although the Oxytocin-REMD-S simulation did not converge (Fig. 6a), it did produce a similar ensemble at 300K as the other replica-exchange-based simulations (Fig. 6g). That is, through similarity analysis in Fig. 6g, the four replica-exchange-based simulations (REMD-{L,S} and ResREMD-{L,S}) sampled similar ensembles at 300K in 200 ns, as indicated by the mutually low JSDs and high torsional similarity values (See Section 2.6). The mutually low JSD and high torsional similarity values were also observed for the four replica-exchange-based simulations for cyclo-(CGPRRARSG) in Fig. 6h. For cyclo-(CGPRRARSG), the Res-REMD simulations do not show significant acceleration of convergence over the REMD simulations (Fig. 6b). These similarity measures also indicate that replica-exchange-based simulations with a “small” temperature range (REMD-S or ResREMD-S) are sufficient to generate correct and reproducible ensembles at 300K for oxytocin and cyclo-(CGPRRARSG).

Consistent with the entropy convergence analysis, the ensembles generated by conventional MD at 300K are different from the replica-exchange-based simulations by similarity analysis: By comparing the torsional state probabilities between pairs of ensembles (blue dots in each scatter plot in the lower triangle of Figs. 6g,h), we found that the conventional MD simulations did not sample all the major component states for oxytocin and cyclo-(CGPRRARSG). For oxytocin, REMD or Res-REMD ensembles consisted of four major torsional states (the 4 points with significant occupation probability in Fig. 6g), but conventional MD can only sample two of them within the simulation time (bottom row in Fig. 6g). Similarly, for cyclo-(CGPRRARSG), the two states with the highest occupation probability in the conventional MD ensemble have low probabilities in the REMD or Res-REMD simulations, supporting the notion that they correspond to kinetic traps (last row in Fig. 6h).

#### 3.3.3 Res-REMD accelerates ensemble generation over REMD for cyclo-(CGHHQKLVG)

For cyclo-(CGHHQKLVG), the REMD simulation with a higher upper bound temperature (CGHHQKLVG-REMD-L) did not converge within the simulation time (Fig. 6c). The torsional entropy has a slowly increasing trend, approaching the entropy of the ensemble generated by Res-REMD with the same temperature range (i.e. by CGHHQKLVG-ResREMD-L). Despite the flattening of the curve corresponding to the torsional entropy of the CGHHQKLVG-REMD-S simulation, it is unlikely to have converged, as it must eventually reach the higher torsional entropy of the CGHHQKLVG-ResREMD-L,exL simulations. The 300K ensemble in the CGHHQKLVG-ResREMD-S simulation on the other hand shows signs of possible convergence, however, the ensemble appears to be converging to an incorrect ensemble with lower torsional entropy than that obtained from the CGHHQKLVG-ResREMD-{L,exL} simulations. We discuss this important problem below in Section 3.3.4. The increasing trend is also observed when we compare the torsional similarity between accumulating portions of the CGHHQKLVG-REMD-L ensemble at 300K and the full CGHHQKLVG-ResREMD-L ensemble at 300K (Fig. 7a). This indicates that the ensemble composition of non-converging CGHHQKLVG-REMD-L was approaching that of the full trajectory of converged CGHHQKLVG-ResREMD-L.

**Figure 7:**
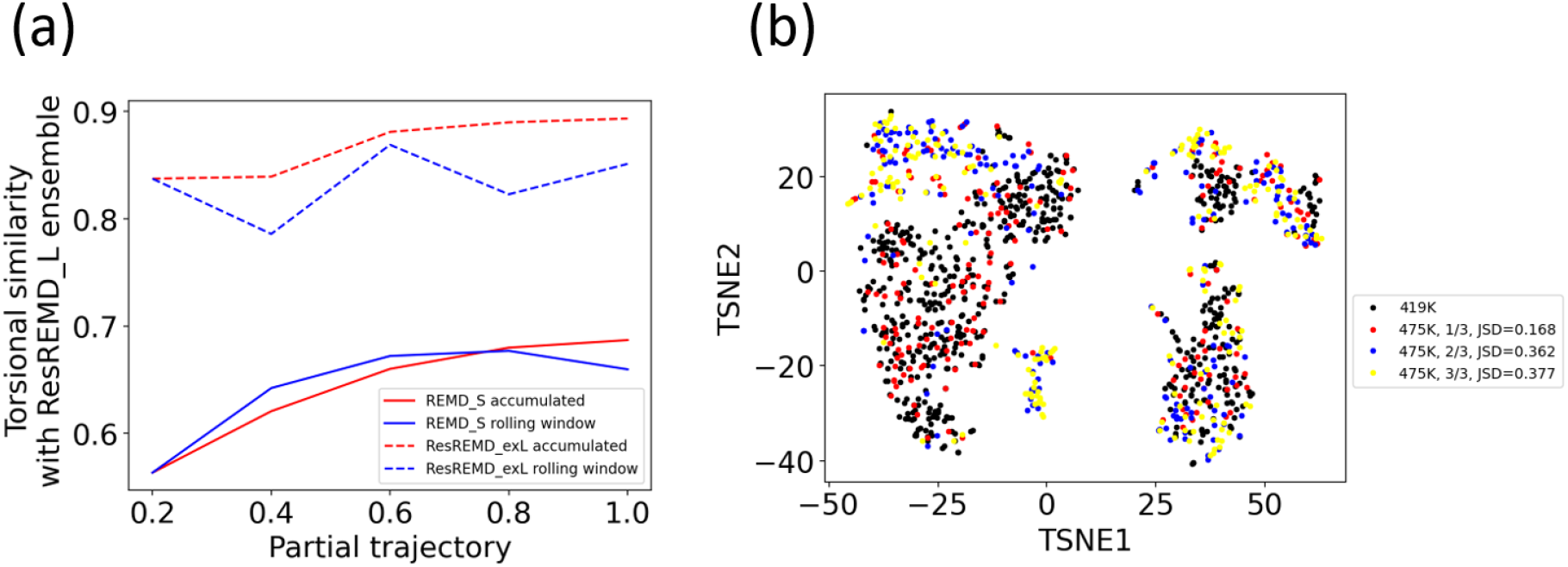
Detailed convergence analysis for cyclo-(CGHHQKLVG) simulations. (a) Subsets of ensembles of REMD-S (solid lines) or ResREMD-exL (dotted lines) at 300K were compared with the full ensemble of ResREMD-L ensemble, again at 300K, using torsional similarities (see Section 2.6.1). Red solid line: The accumulated portions of trajectories of REMD-L at 300K are compared with the full trajectory of ResREMD-L at 300K. Blue solid line: 20% of the trajectory collected in a rolling window for the REMD-L simulation at 300K is compared with the full trajectory of the ResREMD-L simulation at 300K. The dotted lines correspond to the counterparts of the solid lines, but for the ResREMD-exL simulation. (b) The structural phase space of HHQKLV segment in the reservoir ensembles are projected to 2 dimensions using t-SNE (see Section 2.8.1). The configurations in 419K reservoir are in black dots. For the 475K reservoir, the configurations generated from the first 1*/*3 of the trajectory are in red, the second 1*/*3 configurations are in blue, and the last 1*/*3 configurations are in yellow. The extent of overlap between subsets of the 475K reservoir and the 419K reservoir was quantified by JSD which was illustrated in the legend (see Fig. S3 for detail).

By entropy and ensemble similarity analysis above, one might directly conclude that CGHHQKLVG-ResREMD-L generated a more accurate ensemble at 300K than CGHHQKLVG-REMD-L did. However, the 300K ensemble generated by CGHHQKLVG-ResREMD-L is still different from the other reservior-based simulation, CGHHQKLVG-ResREMD-S (Fig. 6i). Thus, we performed an additional Res-REMD simulation, CGHHQKLVG-ResREMD-exL, with a higher reservoir temperature at 537K, to examine whether a similar 300K ensemble is reproduced. The similarity analysis in Fig. 6i shows that CGHHQKLVG-ResREMD-exL and CGHHQKLVG-ResREMD-L generated the most similar ensemble at 300K among the five ensembles (JSD=0.10, and torsional similarity=0.893). The first 20% trajectory of the 300K ensemble of CGHHQKLVG-ResREMD-exL already achieved approximately 0.83 torsional similarity to the full trajectory of the 300K ensemble of CGHHQKLVG-ResREMD-L (Fig. 7a). The above similarity analysis supports the assumption that CGHHQKLVG-ResREMD-L produced an ensemble that is reliable and reproducible by another independent Res-REMD simulation. With the additional Res-REMD simulation (CGHHQKLVG-ResREMD-exL), we confirmed that Res-REMD can accelerate the ensemble generation over REMD for cyclo-(CGHHQKLVG).

#### 3.3.4 Reservoir temperature should be high enough to prevent insufficient sampling in the reservoir generation process

We have illustrated in detail in Section 3.2.3 that an accurate reservoir composition is paramount to the generation of a reliable ensemble by Res-REMD. Thus, it’s not surprising that the inability of CGHHQKLVG-ResREMD-S to generate a correct ensemble at 300K is due to the coupling to a non-equilibrium reservoir at 419K: The torsional entropy of the 419K reservoir (Fig. 6f) converged to a lower value than the correctly converged 300K ensemble as obtained from either the CGHHQKLVG-ResREMD-L or CGHHQKLVG-ResREMD-exL simulations (Fig. 6c), which indicates that insufficient sampling to obtain an equilibrium ensemble in the 419K reservoir results in convergence to a non-equilibrium (incorrect) ensemble at 300K. Similar effects have been observed in Kasavajhala et al.^86^

To further investigate the composition of the reservoir when accumulating states to generate an equilibrium ensemble in the reservoir, the conformational states of conventional MD at 419K and 475K were evenly sampled and projected on 2 dimensional phase space by t-SNE analysis^97^ (see Section 2.8.1) (Fig. 7b). The kernel density estimation of the 419K reservoir (black dots) greatly overlaps with that of the beginning one-third of the trajectory of the 475K reservoir (red dots), with a JSD=0.168 (Fig. S3), but it becomes more dissimilar as the trajectory accumulates conformations–the middle and last one-third of the trajectory of the 475K reservoir (blue and yellow dots) have JSD=0.362 and JSD=0.377, respectively with the 419K reservoir ensemble. This suggests that the reservoir at 419K was trapped in the local free energy basin. The incorrect 300K ensemble generated by CGHHQKLVG-REMD-S, which spans the same temperature range as CGHHQKLVG-ResREMD-S, may also be due to insufficient sampling at high temperature replicas.

Interestingly, although the reservoir at 419K and 475K have lower torsional similarity (0.690 in Fig. 6f inset) than the other cyclic peptides (Fig. 6d,e inset), they have the same first and second most-populated torsional states (top right blue dots in Fig. 6f inset). Therefore, the problems arising from a non-equilibrium reservoir in the CGHHQKLVG-ResREMD-S simulations stem from incomplete sampling of low-populated states at 419K.

#### 3.3.5 Comparison with NMR experiment for oxytocin

Chemical shift and J-coupling values for oxytocin were compared between experimentally measured values in Ohno et al.,^82^ and computed values in this work. Computed chemical shifts were determined using SHIFTX2,^103^ and reported as averages across the entire ensemble. We present secondary chemical shifts, in which residue-specific random coil chemical shifts^104,105^ have been subtracted from both the computed and experimental data. The J-couplings between the HN and H*α* atoms, *J*_*H*_*N Hα*, were calculated for all residues using the Karplus equation,^106^ as implemented in the GROMACS *chi* module. The 300K ensembles generated by Oxytocin-REMD-L, Oxytocin-ResREMD-L, and conventional MD were included in the comparison. The experimentally determined NMR structure (PDB ID: 2MGO) was used as the initial condition in all simulations. We also compare our results with a previous simulation study by Yedvabny et al. ^32^ in which oxytocin conformational ensembles were acquired by Res-REMD in AMBER^37^ with ff99SB-ildn^107^ force field and TIP4P-Ew water model.^108^

The computed 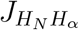 coupling and the secondary chemical shift for C_*α*_, C_*β*_, H_*α*_, H_N_, C, and N atoms were compared with the experimental values in Fig. 8a. The J-couplings as well as the secondary chemical shifts computed from REMD and Res-REMD are close to each other, and they sometimes deviate from the values computed from conventional MD ensemble (correlations between the ensembles are *r*(ResREMD,REMD) = 0.99, *r*(ResREMD,MD) = 0.92, *r*(REMD,MD) = 0.90, see Figs. S3a-c), which is consistent with the similarity analysis wherein the MD ensemble was the most distinct from all other ensembles (Fig 6g). We generated scatter plots for computed *vs*. experimental secondary chemical shifts of C_*α*_, C_*β*_, H_*α*_, and HN atoms for each ensemble (Fig. 8b). It can be observed that the ResREMD-L ensemble has the highest correlation coefficient (*r* = 0.57 in Fig. 8b), which is comparable to the agreement seen previously by Yedvabny et al.^32^ The improvement of Res-REMD over the ensembles obtained by the other methods is not statistically significant however.

**Figure 8:**
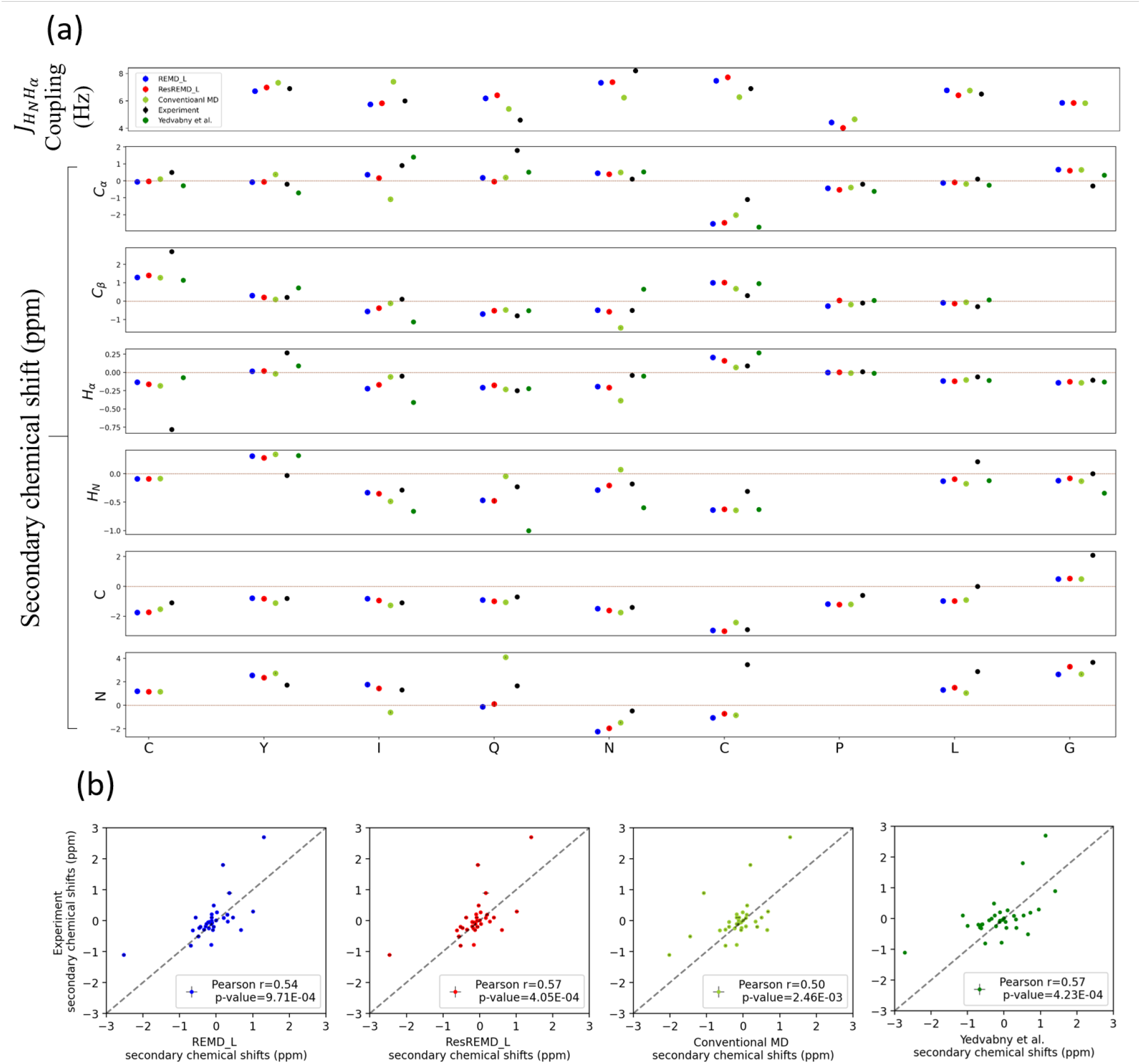
(a) Comparison of the computed and experimental oxytocin *J*_*H*_*N Hα* coupling, as well as the secondary chemical shifts, for C_*α*_, C_*β*_, H_*α*_, H_N_, C, and N. (b) The secondary chemical shifts of C_*α*_, C_*β*_, H_*α*_, H_N_ are pooled and compared between the individual ensemble and the experiment. From left to right are ensemble generated by Oxytocin-REMD-L, Oxytocin-ResREMD-L, conventional MD, and the Yedvabny et al.’s^32^ ensemble at 300K. The vertical error bars in (a) and horizontal error bars in (b) are reported as the standard deviation of the mean. The standard error in the mean is generally smaller than the size of the data points rendered.

Perhaps surprisingly, the conventional MD simulation also exhibited a moderately high correlation with the experimental data (*r* = 0.50). This might be due to the insensitivity of the chemical shift, as an ensemble-averaged measure, to discern sub-ensemble information. Previous studies on A*β* peptides have also found that chemical shifts did not effectively distinguish between different IDP ensembles.^109^ Notably, the correlation between the chemical shift values of our ResREMD-L ensemble and Yedvabny et al.’s^32^ Res-REMD ensemble is *r* = 0.88 (Figs. S3e). This agreement again supports the validity of our ResREMD implementation in GROMACS.

### 3.4 Which cyclic peptide sequences would be predicted to benefit from Res-REMD over REMD?

We found that the degree of sampling acceleration of ResREMD over REMD was not a generic property, but rather was sequence-specific for cyclic peptides. That is, ResREMD simulations only showed faster convergence than REMD for the cyclo-(CGHHQKLVG), but not for the other cyclic peptides. This raises a question: Which cyclic peptide sequences would be predicted to benefit from ResREMD *vs*. REMD by showing accelerated convergence? To answer this question, we performed a convergence analysis of the reservoir generation process, using torsional angle auto-correlation analysis and time-lagged Independent Component Analysis (tICA) (Fig. 9).

**Figure 9:**
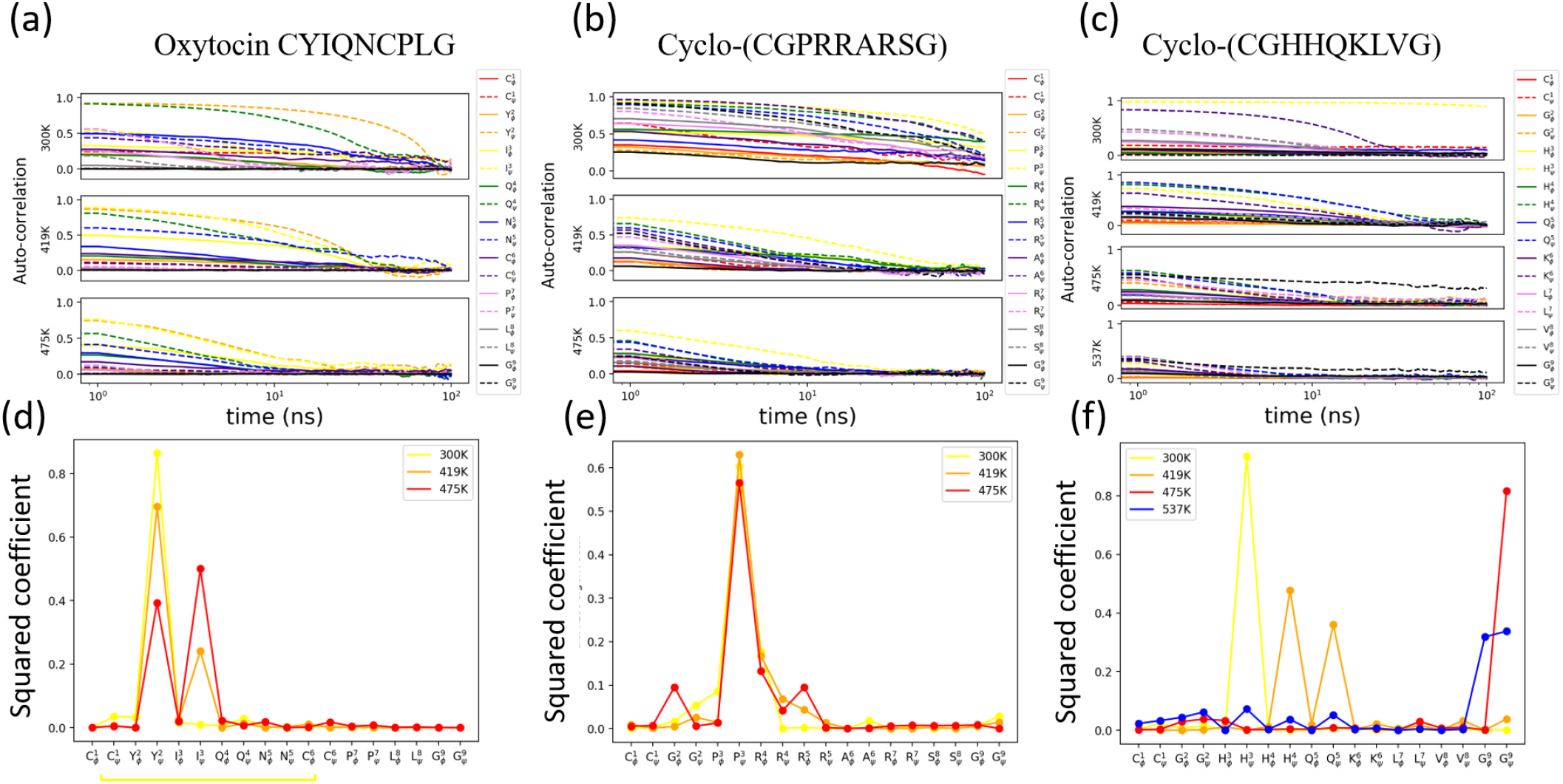
(a-c) The torsional angle auto-correlation functions were calculated for all the conventional MD simulations at temperatures 300K, and the reservoir temperatures 419K, 475K, and 537K for Cyclo-(CGHHQKLVG). (d-f) The squared coefficient of the first tICA eigenvector that corresponds to each system in (a-c), which identifies the slowest converging reaction coordinate. The yellow underline in (d) shows the position of the disulfide bond determining the cyclic portion of the peptide.

Most of the conventional MD simulations at reservoir temperatures 419K, 475K, and 537K have auto-correlations of cosine of torsional angle that decrease with increasing temperature, and decrease to zero within 100 nanoseconds (Fig. 9a-c). An exception appears for the conventional MD at 475K for cyclo-(CGHHQKLVG): the auto-correlation of glycine *ψ* angle at position 9 (G_*ψ*_^9^) maintains a value of *≈* 0.3 up to 100 ns. This corresponds to the upper bound temperature of the only non-converging REMD simulation that did not acquire a correct ensemble, CGHHQKLVG-REMD-L.

As a comparison, auto-correlation analysis was also performed on conventional MD simulations at 300K. The auto-correlation functions at 300K generally decay more slowly than those at reservoir temperatures. Interestingly, oxytocin at 300K shows less auto-correlation than the other cyclic peptides despite the fact that it has a smaller ring.

The slowest reaction coordinate of a linear combination of the cosines of torsional angles was determined from tICA (see Section 2.8.2). The slow-converging torsional angles are those with high squared tICA coefficients (y-axes in Fig. 9d-f). The slow-converging torsional angles were located at select residues constrained within the disulfide bond for oxytocin (Fig. 9d), and the proline for cyclo-(CGPRRARSG) (Fig. 9e). For cyclo-(CGHHQKLVG) (Fig. 9f), the slow-converging torsional angles were located at the epitope (HHQKLV) region at 300K and 419K, but at higher temperatures 475K and 537K, the flanking glycines were the slowest to decorrelate. This finding is consistent with the auto-correlation analysis in Fig. 9c showing that G_*ψ*_^9^ is a slowly converging torsional angle for cyclo-(CGHHQKLVG) at 475K.

We extracted the torsional angles G_*ϕ*_^9^ and G_*ψ*_^9^, from Glycine 9 in the three cyclic peptides analyzed here, at temperatures 419K and 475K, and constructed free energy surface along the Ramachandran space using the histogram method (Fig. S2). Since G^9^ of oxytocin is located at the flanked linear region, it was less constrained than G^9^ in either cyclo-(CGPRRARSG) or cyclo-(CGHHQKLVG). Moreover, G^9^ in cyclo-(CGHHQKLVG) was more constrained than G^9^ in cyclo-(CGPRRARSG). Thus, the constraining effects of cyclization on the Ramachandran space of glycines are sequence specific.

The above analysis gives a method for determining whether it is beneficial to apply Res-REMD rather than REMD. That is, before running REMD or Res-REMD, a conventional MD at the proposed upper-bound temperature can be run first. If the auto-correlations of all cosine of torsional angles decay to zero in, say, 100 nanoseconds, a relatively rapid convergence of the REMD simulation can be ensured by the efficient sampling of the highest-temperature replica in REMD. However, in the event of slow convergence of the highest temperature replica, Res-REMD would be recommended. One limitation of this method is that it cannot determine if the selected temperature is sufficiently high to obtain correct convergence. For example, an incorrectly converged conventional MD simulation with fast-decayed auto-correlation was illustrated by cyclo-(CGHHQKLVG) at 419K (Fig. 9c).

## 4 Conclusion

A reliable Res-REMD simulation tool based on GROMACS 4.6.7 was developed in this study, and tested on five model systems: Alanine dipeptide, Trpzip2, and three cyclic peptides. The alanine dipeptide simulations demonstrated the validity of Res-REMD implementation in GROMACS, and the Trpzip2 simulations demonstrated the efficiency of Res-REMD over REMD in explicit solvent. The sampling acceleration of Res-REMD over REMD for cyclic peptide was found to be sequence specific: Among three cyclic peptides, Res-REMD only showed efficiency gain over REMD in cyclo-(CGHHQKLVG), while the other two cyclic peptides, oxytocin and cyclo-(CGPRRARSG), did not gain sampling acceleration. We found that REMD was unable to acquire sufficient sampling for cyclo-(CGHHQKLVG) due to large torsional angle auto-correlations even at high temperatures. Res-REMD solves this problem by substituting the highest-temperature replica with a pre-sampled converged reservoir. Using the auto-correlation function for torsional angles, one could also determine if Res-REMD should be employed for the generation of cyclic peptide ensembles rather than REMD. We found that generation of a converged equilibrium reservoir was essential for convergence of the simulation to the correct equilibrium ensemble at 300K. Otherwise Res-REMD would only accellerate convergence to a biased (non-equilibrium) ensemble.

The cyclic peptides chosen in this study have several potential applications. The cyclized A*β* segment, cyclo-(CGHHQKLVG), as well as the cyclized furin cleavage site of SARS-CoV-2 spike, cyclo-(CGPRRARSG), could be used as “glycindel” conformationally scaffolded epitopes.^43^ They can be utilized to generate antibodies through active immunization in mice or rabbits.^41,42^ Oxytocin (CYIQNCPLG) was useful as a reference to compare with the NMR experimental data, as well as previous computational results. The computed chemical shifts and J-couplings using the Res-REMD-generated 300K ensemble showed the strongest agreement with the experimental numbers.^82^

Currently, Res-REMD has only been available in AMBER software^37^ and only recently has been implemented in Python.^33^ By providing this simulation tool in GROMACS (https://github.com/PlotkinLab/Reservoir-REMD), we hope to make Res-REMD more broadly accessible to a larger group of researchers using molecular dynamics as a tool for discovery and analysis.

## 5 Associated content

### Supporting Information

The Supporting Information is available free of charge at URL.

Figure S1: Additional information on the transition of folded structure throughout the Trpzip2-ResREMD-2 1 simulation

Figure S2: Free energy surfaces as a function of torsional angles G_*ϕ*_^9^ and G_*ψ*_^9^ in the Ramachandran space, for all the conventional MD ensembles of the three cyclic peptides examined in the main text, at temperatures 419K and 475K.

Figure S3: The kernel density estimation probability of the data points for each color in Fig. 7b.

Figure S4: The comparison of the secondary chemical shifts of C_*α*_, C_*β*_, H_*α*_, H_N_ atoms between different pairs of ensembles.

## 6 Author information

### Author Contributions

Conceptualization, S.H., A.A. and S.S.P.; Methodology, S.H. and S.S.P.; Software, S.H.; Formal analysis, S.H. and S.S.P.; Resources, S.S.P; Writing—original draft preparation, S.H.; Writing—review and editing, S.S.P.; Visualization, S.H.; Supervision, S.S.P.; Project administration, S.S.P.; Funding acquisition, S.S.P.

### Notes

The authors declare no competing financial interest.

## 7 Acknowledgement

This research was funded by the Canadian Institute of Health Research Transitional Operating Grant 2682, and by Alberta Innovates Research Team Program Grant PTM13007, Compute Canada Resources for Research Groups RRG 3071, and UBC ARC Sockeye Advanced Research Computing (https://doi.org/10.14288/SOCKEYE,2019). S.H. has received support from an NSERC CREATE–Ecosystem Services, Commercialization, and Entrepreneurship (ECOSCOPE) Scholarship, and Mitacs Accelerate Scholarship. The authors also thank Pranav Garg for her helpful advice on restructuring and improving the quality of an early draft of the manuscript, as well as Gabriel Dall’Alba and Charlie Barclay for their critical reading and suggestions on the manuscript.

## Graphical TOC Entry

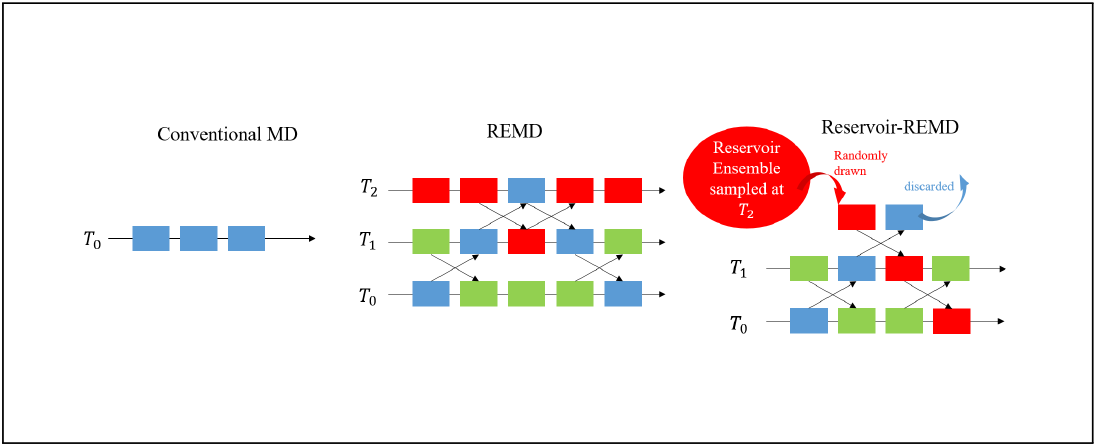

## References

(1) Frauenfelder, H.; Sligar, S. G.; Wolynes, P. G. The energy landscapes and motions of proteins. Science 1991, 254, 1598–1603.

(2) Wales, D. J.; Doye, J. P.; Miller, M. A.; Mortenson, P. N.; Walsh, T. R. Energy landscapes: from clusters to biomolecules. Advances in Chemical Physics 2000, 115, 1–111.

(3) Fisher, C. K.; Stultz, C. M. Constructing ensembles for intrinsically disordered proteins. Current opinion in structural biology 2011, 21, 426–431.

(4) Herschlag, D.; Allred, B. E.; Gowrishankar, S. From static to dynamic: the need for structural ensembles and a predictive model of RNA folding and function. Current opinion in structural biology 2015, 30, 125–133.

(5) Bonomi, M.; Heller, G. T.; Camilloni, C.; Vendruscolo, M. Principles of protein structural ensemble determination. Current opinion in structural biology 2017, 42, 106–116.

(6) Onuchic, J. N.; Luthey-Schulten, Z.; Wolynes, P. G. Theory of Protein Folding: The Energy Landscape Perspective. Annual Review of Physical Chemistry 1997, 48, 545– 600.

(7) Nymeyer, H.; García, A. E.; Onuchic, J. N. Folding funnels and frustration in off-lattice minimalist protein landscapes. Proceedings of the National Academy of Sciences 1998, 95, 5921–5928.

(8) Shea, J.-E.; Onuchic, J. N.; Brooks III, C. L. Exploring the origins of topological frustration: Design of a minimally frustrated model of fragment B of protein A. Proceedings of the National Academy of Sciences 1999, 96, 12512–12517.

(9) Clementi, C.; Nymeyer, H.; Onuchic, J. N. Topological and energetic factors: what determines the structural details of the transition state ensemble and “en-route” intermediates for protein folding? An investigation for small globular proteins. Journal of molecular biology 2000, 298, 937–953.

(10) Cheung, M. S.; García, A. E.; Onuchic, J. N. Protein folding mediated by solvation: Water expulsion and formation of the hydrophobic core occur after the structural collapse. Proceedings of the National Academy of Sciences 2002, 99, 685–690.

(11) Plotkin, S. S.; Onuchic, J. N. Understanding protein folding with energy landscape theory part I: Basic concepts. Quarterly reviews of biophysics 2002, 35, 111–167.

(12) Plotkin, S. S.; Onuchic, J. N. Understanding protein folding with energy landscape theory Part II: Quantitative aspects. Quarterly reviews of biophysics 2002, 35, 205– 286.

(13) Levy, Y.; Onuchic, J. N. Water Mediation in Protein Folding and Molecular Recognition. Annual Review of Biophysics and Biomolecular Structure 2006, 35, 389–415.

(14) Whitford, P. C.; Noel, J. K.; Gosavi, S.; Schug, A.; Sanbonmatsu, K. Y.; Onuchic, J. N. An all-atom structure-based potential for proteins: bridging minimal models with allatom empirical forcefields. Proteins: Structure, Function, and Bioinformatics 2009, 75, 430–441.

(15) Noel, J. K.; Whitford, P. C.; Sanbonmatsu, K. Y.; Onuchic, J. N. N. SMOG@ctbp: simplified deployment of structure-based models in GROMACS. Nucleic Acids Research 2010, 38, W657–W661.

(16) Morcos, F.; Onuchic, J. N. The role of coevolutionary signatures in protein interaction dynamics, complex inference, molecular recognition, and mutational landscapes. Current Opinion in Structural Biology 2019, 56, 179–186.

(17) Hyeon, C.; Onuchic, J. N. A structural perspective on the dynamics of kinesin motors. Biophysical journal 2011, 101, 2749–2759.

(18) Su-lkowska, J. I.; Rawdon, E. J.; Millett, K. C.; Onuchic, J. N.; Stasiak, A. Conservation of complex knotting and slipknotting patterns in proteins. Proceedings of the National Academy of Sciences 2012, 109, E1715–E1723.

(19) Bai, F.; Morcos, F.; Cheng, R. R.; Jiang, H.; Onuchic, J. N. Elucidating the druggable interface of proteinprotein interactions using fragment docking and coevolutionary analysis. Proceedings of the National Academy of Sciences 2016, 113, E8051–E8058.

(20) Lin, X.; Roy, S.; Jolly, M. K.; Bocci, F.; Schafer, N. P.; Tsai, M.-Y.; Chen, Y.; He, Y.; Grishaev, A.; Weninger, K.; Orban, J.; Kulkarni, P.; Rangarajan, G.; Levine, H.; Onuchic, J. N. PAGE4 and Conformational Switching: Insights from Molecular Dynamics Simulations and Implications for Prostate Cancer. Journal of Molecular Biology 2018, 430, 2422–2438.

(21) Cheng, R. R.; Contessoto, V. G.; Aiden, E. L.; Wolynes, P. G.; Di Pierro, M.; Onuchic, J. N. Exploring chromosomal structural heterogeneity across multiple cell lines. Elife 2020, 9, e60312.

(22) Swendsen, R. H.; Wang, J.-S. Replica Monte Carlo Simulation of Spin-Glasses. Phys. Rev. Lett. 1986, 57, 2607–2609.

(23) Sugita, Y.; Okamoto, Y. Replica-exchange molecular dynamics method for protein folding. Chemical physics letters 1999, 314, 141–151.

(24) Nymeyer, H.; Gnanakaran, S.; Garcia, A. E. Atomic simulations of protein folding, using the replica exchange algorithm. 2004, 383, 119–149.

(25) Mitsutake, A.; Sugita, Y.; Okamoto, Y. Generalized-ensemble algorithms for molecular simulations of biopolymers. Peptide Science: Original Research on Biomolecules 2001, 60, 96–123.

(26) García, A. E.; Onuchic, J. N. Folding a protein in a computer: an atomic description of the folding/unfolding of protein A. Proceedings of the National Academy of Sciences 2003, 100, 13898–13903.

(27) Yang, S.; Onuchic, J. N.; García, A. E.; Levine, H. Folding time predictions from allatom replica exchange simulations. Journal of molecular biology 2007, 372, 756–763.

(28) Okur, A.; Roe, D. R.; Cui, G.; Hornak, V.; Simmerling, C. Improving convergence of replica-exchange simulations through coupling to a high-temperature structure reservoir. Journal of chemical theory and computation 2007, 3, 557–568.

(29) Li, H.; Li, G.; Berg, B. A.; Yang, W. Finite reservoir replica exchange to enhance canonical sampling in rugged energy surfaces. The Journal of Chemical Physics 2006, 125, 144902.

(30) Lyman, E.; Ytreberg, F. M.; Zuckerman, D. M. Resolution Exchange Simulation. Phys. Rev. Lett. 2006, 96, 028105.

(31) Ruscio, J. Z.; Fawzi, N. L.; Head-Gordon, T. How hot? Systematic convergence of the replica exchange method using multiple reservoirs. Journal of computational chemistry 2010, 31, 620–627.

(32) Yedvabny, E.; Nerenberg, P. S.; So, C.; Head-Gordon, T. Disordered structural ensembles of vasopressin and oxytocin and their mutants. The Journal of Physical Chemistry B 2015, 119, 896–905.

(33) Fabregat, R.; Fabrizio, A.; Meyer, B.; Hollas, D.; Corminboeuf, C. Hamiltonian-Reservoir Replica Exchange and Machine Learning Potentials for Computational Organic Chemistry. Journal of chemical theory and computation 2020, 16, 3084–3094.

(34) Okur, A.; Miller, B. T.; Joo, K.; Lee, J.; Brooks, B. R. Generating reservoir conformations for replica exchange through the use of the conformational space annealing method. Journal of chemical theory and computation 2013, 9, 1115–1124.

(35) Roitberg, A. E.; Okur, A.; Simmerling, C. Coupling of replica exchange simulations to a non-Boltzmann structure reservoir. The Journal of Physical Chemistry B 2007, 111, 2415–2418.

(36) Henriksen, N. M.; Roe, D. R.; Cheatham III, T. E. Reliable oligonucleotide conformational ensemble generation in explicit solvent for force field assessment using reservoir replica exchange molecular dynamics simulations. The journal of physical chemistry B 2013, 117, 4014–4027.

(37) Case, D.; Aktulga, H.; Belfon, K.; Ben-Shalom, I.; Berryman, J.; Brozell, S.; Cerutti, D.; Cheatham, T.; III,; Cisneros, G.; Cruzeiro, V.; Darden, T.; Duke, R.; Giambasu, G.; Gilson, M.; Gohlke, H.; Goetz, A.; Harris, R.; Izadi, S.; Izmailov, S.; Kasavajhala, K.; Kaymak, M.; King, E.; Kovalenko, A.; Kurtzman, T.; Lee, T.; LeGrand, S.; Li, P.; Lin, C.; Liu, J.; Luchko, T.; Luo, R.; Machado, M.; Man, V.; Manathunga, M.; Merz, K.; Miao, Y.; Mikhailovskii, O.; Monard, G.; Nguyen, H.; O’Hearn, K.; Onufriev, A.; Pan, F.; Pantano, S.; Qi, R.; Rahnamoun, A.; Roe, D.; Roitberg, A.; Sagui, C.; Schott-Verdugo, S.; Shajan, A.; Shen, J.; Simmerling, C.; Skrynnikov, N.; Smith, J.; Swails, J.; Walker, R.; Wang, J.; Wang, J.; Wei, H.; Wolf, R.; Wu, X.; Xiong, Y.; Xue, Y.; York, D.; Zhao, S.; Kollman, P. Amber 2022. 2022,

(38) Mulligan, V. K. The emerging role of computational design in peptide macrocycle drug discovery. Expert Opinion on Drug Discovery 2020, 1–19.

(39) Zorzi, A.; Deyle, K.; Heinis, C. Cyclic peptide therapeutics: past, present and future. Current opinion in chemical biology 2017, 38, 24–29.

(40) Jing, X.; Jin, K. A gold mine for drug discovery: Strategies to develop cyclic peptides into therapies. Medicinal research reviews 2020, 40, 753–810.

(41) Silverman, J. M.; Gibbs, E.; Peng, X.; Martens, K. M.; Balducci, C.; Wang, J.; Yousefi, M.; Cowan, C. M.; Lamour, G.; Louadi, S.; Ban, Y.; Robert, J.; Stukas, S.; Forloni, G.; Hsiung, G.-Y. R.; Plotkin, S. S.; Wellington, C. L.; Cashman, N. R. A Rational Structured Epitope Defines a Distinct Subclass of Toxic Amyloid-beta Oligomers. ACS Chemical Neuroscience 2018, 9, 1591–1606.

(42) Gibbs, E.; Silverman, J. M.; Zhao, B.; Peng, X.; Wang, J.; Wellington, C. L.; Mackenzie, I. R.; Plotkin, S. S.; Kaplan, J. M.; Cashman, N. R. A rationally designed humanized antibody selective for amyloid beta oligomers in Alzheimer’s disease. Scientific reports 2019, 9, 1–14.

(43) Hsueh, S. C. C.; Aina, A.; Roman, A. Y.; Cashman, N. R.; Peng, X.; Plotkin, S. S. Optimizing Epitope Conformational Ensembles Using α-Synuclein Cyclic Peptide “Glycindel” Scaffolds: A Customized Immunogen Method for Generating Oligomer-Selective Antibodies for Parkinson’s Disease. ACS Chemical Neuroscience 2022, https://doi.org/10.1021/acschemneuro.1c00567.

(44) Driggers, E. M.; Hale, S. P.; Lee, J.; Terrett, N. K. The exploration of macrocycles for drug discovery—an underexploited structural class. Nature Reviews Drug Discovery 2008, 7, 608–624.

(45) Geng, H.; Jiang, F.; Wu, Y.-D. Accurate structure prediction and conformational analysis of cyclic peptides with residue-specific force fields. The journal of physical chemistry letters 2016, 7, 1805–1810.

(46) Gavenonis, J.; Sheneman, B. A.; Siegert, T. R.; Eshelman, M. R.; Kritzer, J. A. Comprehensive analysis of loops at protein-protein interfaces for macrocycle design. Nature chemical biology 2014, 10, 716–722.

(47) Yu, H.; Lin, Y.-S. Toward structure prediction of cyclic peptides. Physical Chemistry Chemical Physics 2015, 17, 4210–4219.

(48) Slough, D. P.; McHugh, S. M.; Cummings, A. E.; Dai, P.; Pentelute, B. L.; Kritzer, J. A.; Lin, Y.-S. Designing well-structured cyclic pentapeptides based on sequence–structure relationships. The Journal of Physical Chemistry B 2018, 122, 3908–3919.

(49) Thevenet, P.; Shen, Y.; Maupetit, J.; Guyon, F.; Derreumaux, P.; Tuffery, P. PEP-FOLD: an updated de novo structure prediction server for both linear and disulfide bonded cyclic peptides. Nucleic acids research 2012, 40, W288–W293.

(50) Hosseinzadeh, P.; Bhardwaj, G.; Mulligan, V. K.; Shortridge, M. D.; Craven, T. W.; Pardo-Avila, F.; Rettie, S. A.; Kim, D. E.; Silva, D.-A.; Ibrahim, Y. M.; Webb, I. K.; Cort, J. R.; Adkins, J. N.; Varani, G.; Baker, D. Comprehensive computational design of ordered peptide macrocycles. Science 2017, 358, 1461–1466.

(51) Mulligan, V. K.; Workman, S.; Sun, T.; Rettie, S.; Li, X.; Worrall, L. J.; Craven, T. W.; King, D. T.; Hosseinzadeh, P.; Watkins, A. M.; Renfrew, P. D.; Guffy, S.; Labonte, J. W.; Moretti, R.; Bonneau, R.; Strynadka, N. C. J.; Baker, D. Computationally designed peptide macrocycle inhibitors of New Delhi metallo-betalactamase 1. Proceedings of the National Academy of Sciences 2021, 118, e2012800118.

(52) Guardiola, S.; Varese, M.; Roig, X.; Sánchez-Navarro, M.; García, J.; Giralt, E. Target-templated de novo design of macrocyclic d-/l-peptides: discovery of drug-like inhibitors of PD-1. Chemical Science 2021, 12, 5164–5170.

(53) Peraro, L.; Kritzer, J. A. Emerging methods and design principles for cell-penetrant peptides. Angewandte Chemie International Edition 2018, 57, 11868–11881.

(54) Witek, J.; Keller, B. G.; Blatter, M.; Meissner, A.; Wagner, T.; Riniker, S. Kinetic models of cyclosporin A in polar and apolar environments reveal multiple congruent conformational states. Journal of chemical information and modeling 2016, 56, 1547– 1562.

(55) Witek, J.; Mühlbauer, M.; Keller, B. G.; Blatter, M.; Meissner, A.; Wagner, T.; Riniker, S. Interconversion rates between conformational states as rationale for the membrane permeability of cyclosporines. ChemPhysChem 2017, 18, 3309–3314.

(56) Witek, J.; Wang, S.; Schroeder, B.; Lingwood, R.; Dounas, A.; Roth, H.-J.; Fouché, M.; Blatter, M.; Lemke, O.; Keller, B.; Riniker, S. Rationalization of the membrane permeability differences in a series of analogue cyclic decapeptides. Journal of chemical information and modeling 2018, 59, 294–308.

(57) Haensele, E.; Banting, L.; Whitley, D. C.; Clark, T. Conformation and dynamics of 8-Arg-vasopressin in solution. Journal of molecular modeling 2014, 20, 1–17.

(58) Haensele, E.; Saleh, N.; Read, C. M.; Banting, L.; Whitley, D. C.; Clark, T. Can Simulations and Modeling Decipher NMR Data for Conformational Equilibria? Arginine– Vasopressin. Journal of chemical information and modeling 2016, 56, 1798–1807.

(59) Li, J.; Ehlers, T.; Sutter, J.; Varma-O’Brien, S.; Kirchmair, J. CAESAR: a new conformer generation algorithm based on recursive buildup and local rotational symmetry consideration. Journal of chemical information and modeling 2007, 47, 1923–1932.

(60) Kirchmair, J.; Laggner, C.; Wolber, G.; Langer, T. Comparative analysis of protein-bound ligand conformations with respect to catalyst’s conformational space subsampling algorithms. Journal of chemical information and modeling 2005, 45, 422–430.

(61) Watts, K. S.; Dalal, P.; Tebben, A. J.; Cheney, D. L.; Shelley, J. C. Macrocycle conformational sampling with MacroModel. Journal of chemical information and modeling 2014, 54, 2680–2696.

(62) Labute, P. LowModeMD: implicit low-mode velocity filtering applied to conformational search of macrocycles and protein loops. Journal of chemical information and modeling 2010, 50, 792–800.

(63) Hawkins, P. C.; Skillman, A. G.; Warren, G. L.; Ellingson, B. A.; Stahl, M. T. Conformer generation with OMEGA: algorithm and validation using high quality structures from the Protein Databank and Cambridge Structural Database. Journal of chemical information and modeling 2010, 50, 572–584.

(64) Sindhikara, D.; Spronk, S. A.; Day, T.; Borrelli, K.; Cheney, D. L.; Posy, S. L. Improving accuracy, diversity, and speed with prime macrocycle conformational sampling. Journal of chemical information and modeling 2017, 57, 1881–1894.

(65) Coutsias, E. A.; Lexa, K. W.; Wester, M. J.; Pollock, S. N.; Jacobson, M. P. Exhaustive conformational sampling of complex fused ring macrocycles using inverse kinematics. Journal of chemical theory and computation 2016, 12, 4674–4687.

(66) Miao, J.; Descoteaux, M. L.; Lin, Y.-S. Structure prediction of cyclic peptides by molecular dynamics+ machine learning. Chemical science 2021, 12, 14927–14936.

(67) McHugh, S. M.; Rogers, J. R.; Yu, H.; Lin, Y.-S. Insights into how cyclic peptides switch conformations. Journal of chemical theory and computation 2016, 12, 2480– 2488.

(68) Kamenik, A. S.; Lessel, U.; Fuchs, J. E.; Fox, T.; Liedl, K. R. Peptidic macrocycles-conformational sampling and thermodynamic characterization. Journal of chemical information and modeling 2018, 58, 982–992.

(69) Ono, S.; Naylor, M. R.; Townsend, C. E.; Okumura, C.; Okada, O.; Lokey, R. S. Conformation and permeability: cyclic hexapeptide diastereomers. Journal of chemical information and modeling 2019, 59, 2952–2963.

(70) Cochran, A. G.; Skelton, N. J.; Starovasnik, M. A. Tryptophan zippers: Stable, monomeric β-hairpins. Proceedings of the National Academy of Sciences 2001, 98, 5578–5583.

(71) Koehbach, J.; O’Brien, M.; Muttenthaler, M.; Miazzo, M.; Akcan, M.; Elliott, A. G.; Daly, N. L.; Harvey, P. J.; Arrowsmith, S.; Gunasekera, S.; Smith, T. J.; Wray, S.; Göransson, U.; Dawson, P. E.; Craik, D. J.; Freissmuth, M.; Gruber, C. W. Oxytocic plant cyclotides as templates for peptide G protein-coupled receptor ligand design. Proceedings of the National Academy of Sciences 2013, 110, 21183–21188.

(72) Choules, M. P.; Bisson, J.; Gao, W.; Lankin, D. C.; McAlpine, J. B.; Niemitz, M.; Jaki, B. U.; Franzblau, S. G.; Pauli, G. F. Quality control of therapeutic peptides by 1H NMR HiFSA sequencing. The Journal of organic chemistry 2019, 84, 3055–3073.

(73) Cashman, N. R.; Plotkin, S. S. N-terminal epitopes in beta-amyloid and conformationally selective antibodies thereof. 2019; European Patent Office EP3374379A4.

(74) Peng, X.; Cashman, N. R.; Plotkin, S. S. Prediction of Misfolding-Specific Epitopes in SOD1 Using Collective Coordinates. The Journal of Physical Chemistry B 2018, 122, 11662–11676.

(75) Johnson, B. A.; Xie, X.; Bailey, A. L.; Kalveram, B.; Lokugamage, K. G.; Muruato, A.; Zou, J.; Zhang, X.; Juelich, T.; Smith, J. K.; Zhang, L.; Bopp, N.; Schindewolf, C.; Vu, M.; Vanderheiden, A.; Winkler, E. S.; Swetnam, D.; Plante, J. A.; Aguilar, P.; Plante, K. S.; Popov, V.; Lee, B.; Weaver, S. C.; Suthar, M. S.; Routh, A. L.; Ren, P.; Ku, Z.; An, Z.; Debbink, K.; Diamond, M. S.; Shi, P.-Y.; Freiberg, A. N.; Menachery, V. D. Loss of furin cleavage site attenuates SARS-CoV-2 pathogenesis. Nature 2021, 591, 293–299.

(76) Spelios, M. G.; Capanelli, J. M.; Li, A. W. A novel antibody against the furin cleavage site of SARS-CoV-2 spike protein: Effects on proteolytic cleavage and ACE2 binding. Immunology Letters 2022, 242, 1–7.

(77) Saito, A.; Irie, T.; Suzuki, R.; Maemura, T.; Nasser, H.; Uriu, K.; Kosugi, Y.; Shirakawa, K.; Sadamasu, K.; Kimura, I.; Ito, J.; Wu, J.; Iwatsuki-Horimoto, K.; Ito, M.; Yamayoshi, S.; Loeber, S.; Tsuda, M.; Wang, L.; Ozono, S.; Butlertanaka, E. P.; Tanaka, Y. L.; Shimizu, R.; Shimizu, K.; Yoshimatsu, K.; Kawabata, R.; Sakaguchi, T.; Tokunaga, K.; Yoshida, I.; Asakura, H.; Nagashima, M.; Kazuma, Y.; Nomura, R.; Horisawa, Y.; Yoshimura, K.; Takaori-Kondo, A.; Imai, M.; Chiba, M.; Furihata, H.; Hasebe, H.; Kitazato, K.; Kubo, H.; Misawa, N.; Morizako, N.; Noda, K.; Oide, A.; Suganami, M.; Takahashi, M.; Tsushima, K.; Yokoyama, M.; Yuan, Y.; Tanaka, S.; Nakagawa, S.; Ikeda, T.; Fukuhara, T.; Kawaoka, Y.; Sato, K.; to Phenotype Japan (G2P-Japan) Consortium, T. G. Enhanced fusogenicity and pathogenicity of SARS-CoV-2 Delta P681R mutation. Nature 2022, 602, 300–306.

(78) Mohammad, A.; Abubaker, J.; Al-Mulla, F. Structural modelling of SARS-CoV-2 alpha variant (B.1.1.7) suggests enhanced furin binding and infectivity. Virus Research 2021, 303, 198522.

(79) Bonomi, M.; Branduardi, D.; Bussi, G.; Camilloni, C.; Provasi, D.; Raiteri, P.; Donadio, D.; Marinelli, F.; Pietrucci, F.; Broglia, R. A.; Parrinello, M. PLUMED: A portable plugin for free-energy calculations with molecular dynamics. Computer Physics Communications 2009, 180, 1961–1972.

(80) Lu, J.-X.; Qiang, W.; Yau, W.-M.; Schwieters, C. D.; Meredith, S. C.; Tycko, R. Molecular structure of β-amyloid fibrils in Alzheimer’s disease brain tissue. Cell 2013, 154, 1257–1268.

(81) Deepmind, Computational predictions of protein structures associated with COVID-19. https://deepmind.com/research/open-source/computational-predictions-of-protein-structures-associated-with-COVID-19 2020,

(82) Ohno, A.; Kawasaki, N.; Fukuhara, K.; Okuda, H.; Yamaguchi, T. Complete NMR analysis of oxytocin in phosphate buffer. Magnetic Resonance in Chemistry 2010, 48, 168–172.

(83) Huang, J.; Rauscher, S.; Nawrocki, G.; Ran, T.; Feig, M.; De Groot, B. L.; Grubmüller, H.; MacKerell, A. D. CHARMM36m: an improved force field for folded and intrinsically disordered proteins. Nature methods 2017, 14, 71–73.

(84) Jorgensen, W. L.; Chandrasekhar, J.; Madura, J. D.; Impey, R. W.; Klein, M. L. Comparison of simple potential functions for simulating liquid water. The Journal of chemical physics 1983, 79, 926–935.

(85) Patriksson, A.; van der Spoel, D. A temperature predictor for parallel tempering simulations. Physical Chemistry Chemical Physics 2008, 10, 2073–2077.

(86) Kasavajhala, K.; Lam, K.; Simmerling, C. Exploring Protocols to Build Reservoirs to Accelerate Temperature Replica Exchange MD Simulations. Journal of Chemical Theory and Computation 2020, 16, 7776–7799.

(87) Marcos, E.; Chidyausiku, T. M.; McShan, A. C.; Evangelidis, T.; Nerli, S.; Carter, L.; Nivón, L. G.; Davis, A.; Oberdorfer, G.; Tripsianes, K.; Sgourakis, N. G.; Baker, D. De novo design of a non-local β-sheet protein with high stability and accuracy. Nature Structural & Molecular Biology 2018, 25, 1028–1034.

(88) Begon, M.; Harper, J.; Townsend, C. Ecology: Individuals, Populations and Communities, 3rd edn. Black-well Science. Oxford 1996,

(89) Sibson, R. Information radius. Zeitschrift für Wahrscheinlichkeitstheorie und verwandte Gebiete 1969, 14, 149–160.

(90) Lin, J. Divergence measures based on the Shannon entropy. IEEE Transactions on Information theory 1991, 37, 145–151.

(91) Lindorff-Larsen, K.; Ferkinghoff-Borg, J. Similarity Measures for Protein Ensembles. PLOS ONE 2009, 4, 1–13.

(92) Tiberti, M.; Papaleo, E.; Bengtsen, T.; Boomsma, W.; Lindorff-Larsen, K. ENCORE: Software for Quantitative Ensemble Comparison. PLOS Computational Biology 2015, 11, 1–16.

(93) Kullback, S.; Leibler, R. A. On information and sufficiency. The annals of mathematical statistics 1951, 22, 79–86.

(94) Cover, T.; Thomas, J. Elements of Information Theory. New York: Wiley 1991, 0471062596–9780471062592.

(95) Shirts, M. R.; Chodera, J. D. Statistically optimal analysis of samples from multiple equilibrium states. The Journal of Chemical Physics 2008, 129, 124105.

(96) Hsueh, S. C. C.; Nijland, M.; Peng, X.; Hilton, B.; Plotkin, S. S. First Principles Calculation of Protein-Protein Dimer Affinities of ALS-Associated SOD1 Mutants. Frontiers in Molecular Biosciences 2022, 9, 845013.

(97) Van der Maaten, L.; Hinton, G. Visualizing data using t-SNE. Journal of machine learning research 2008, 9, 2579–2605.

(98) Pérez-Hernández, G.; Paul, F.; Giorgino, T.; De Fabritiis, G.; Noé, F. Identification of slow molecular order parameters for Markov model construction. The Journal of Chemical Physics 2013, 139, 015102.

(99) Schwantes, C. R.; Pande, V. S. Improvements in Markov state model construction reveal many non-native interactions in the folding of NTL9. Journal of chemical theory and computation 2013, 9, 2000–2009.

(100) Chipot, C.; Pohorille, A. Free energy calculations. Springer series in chemical physics 2007, 86, 159–184.

(101) Okur, A.; Strockbine, B.; Hornak, V.; Simmerling, C. Using PC clusters to evaluate the transferability of molecular mechanics force fields for proteins. Journal of computational chemistry 2003, 24, 21–31.

(102) Still, W. C.; Tempczyk, A.; Hawley, R. C.; Hendrickson, T. Semianalytical treatment of solvation for molecular mechanics and dynamics. Journal of the American Chemical Society 1990, 112, 6127–6129.

(103) Han, B.; Liu, Y.; Ginzinger, S. W.; Wishart, D. S. SHIFTX2: significantly improved protein chemical shift prediction. Journal of biomolecular NMR 2011, 50, 43–57.

(104) Schwarzinger, S.; Kroon, G. J.; Foss, T. R.; Wright, P. E.; Dyson, H. J. Random coil chemical shifts in acidic 8 M urea: implementation of random coil shift data in NMRView. Journal of biomolecular NMR 2000, 18, 43–48.

(105) Kjaergaard, M.; Brander, S.; Poulsen, F. M. Random coil chemical shift for intrinsically disordered proteins: effects of temperature and pH. Journal of biomolecular NMR 2011, 49, 139–149.

(106) Karplus, M. Contact electron-spin coupling of nuclear magnetic moments. The Journal of chemical physics 1959, 30, 11–15.

(107) Lindorff-Larsen, K.; Piana, S.; Palmo, K.; Maragakis, P.; Klepeis, J. L.; Dror, R. O.; Shaw, D. E. Improved side-chain torsion potentials for the Amber ff99SB protein force field. Proteins: Structure, Function, and Bioinformatics 2010, 78, 1950–1958.

(108) Horn, H. W.; Swope, W. C.; Pitera, J. W.; Madura, J. D.; Dick, T. J.; Hura, G. L.; Head-Gordon, T. Development of an improved four-site water model for biomolecular simulations: TIP4P-Ew. The Journal of chemical physics 2004, 120, 9665–9678.

(109) Ball, K. A.; Wemmer, D. E.; Head-Gordon, T. Comparison of structure determination methods for intrinsically disordered amyloid-β peptides. The Journal of Physical Chemistry B 2014, 118, 6405–6416.

