## Supporting Information for "Accelerated ensemble generation for cyclic peptides using a Reservoir-REMD implementation in GROMACS"

### Table of contents

Figures


1. S1—Additional information on the transition of folded structure throughout the Trpzip2-ResREMD-2_1 simulation

2. S2—Free energy surfaces as a function of torsional angles ${G_\phi}^9$ and ${G_\psi}^9$ in the Ramachandran space, for all the conventional MD ensembles of the three cyclic peptides examined in the main text, at temperatures 419K and 475K.

3. S3—The kernel density estimation probability of the data points for each color in Fig. 7b.

4. S4—The comparison of the secondary chemical shifts of $C_\alpha$, $C_\beta$, $H_\alpha$, $H_N$ atoms between different pairs of ensembles.

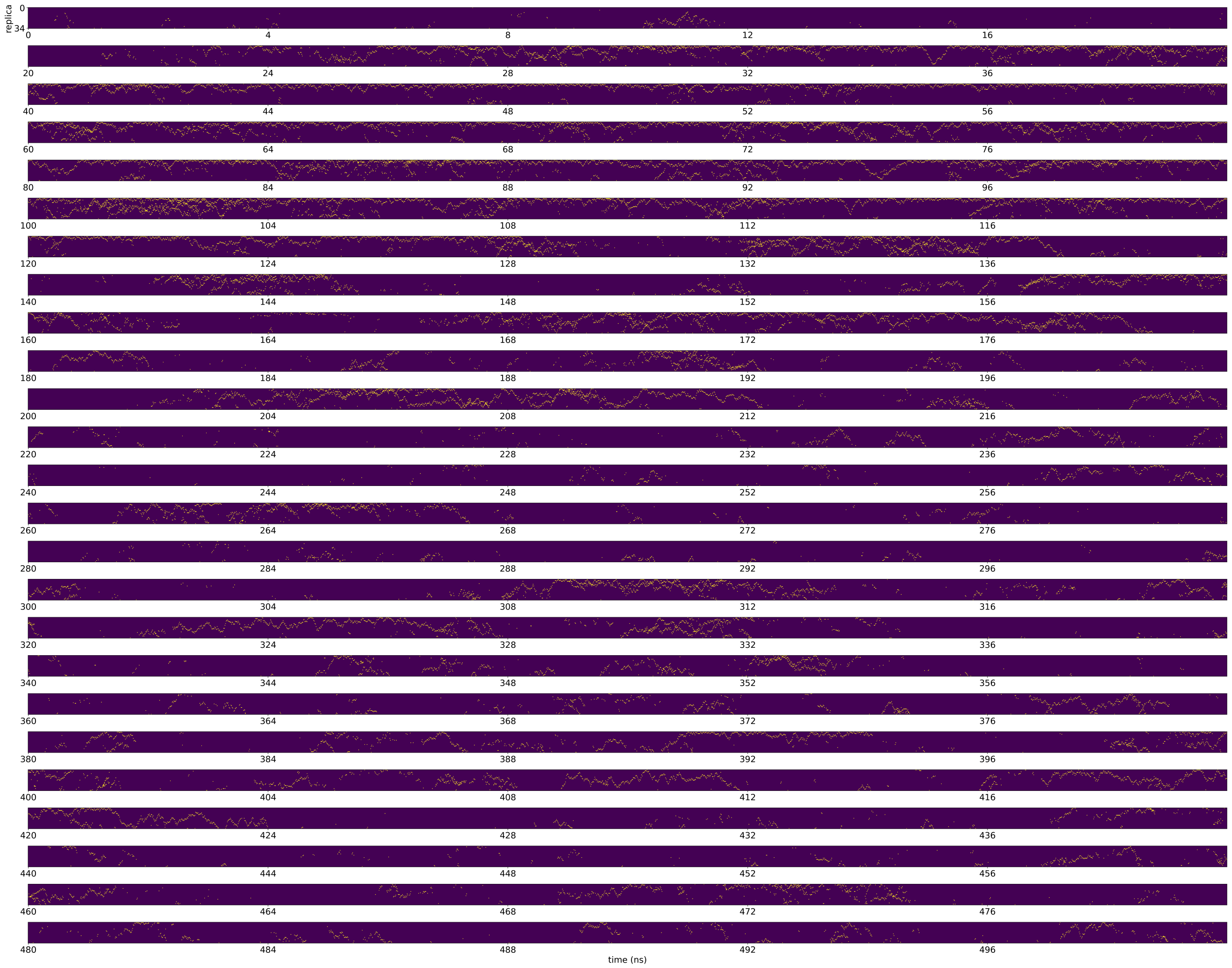


Figure S1: The transition of folded structure throughout the Trpzip2-ResREMD-2_1 simulation. The folded structure with RMSD $< 0.27$ is in yellow, and otherwise blue. The replica index on y-axis from 0 to 35 correspond to the lowest replica temperature to the highest.

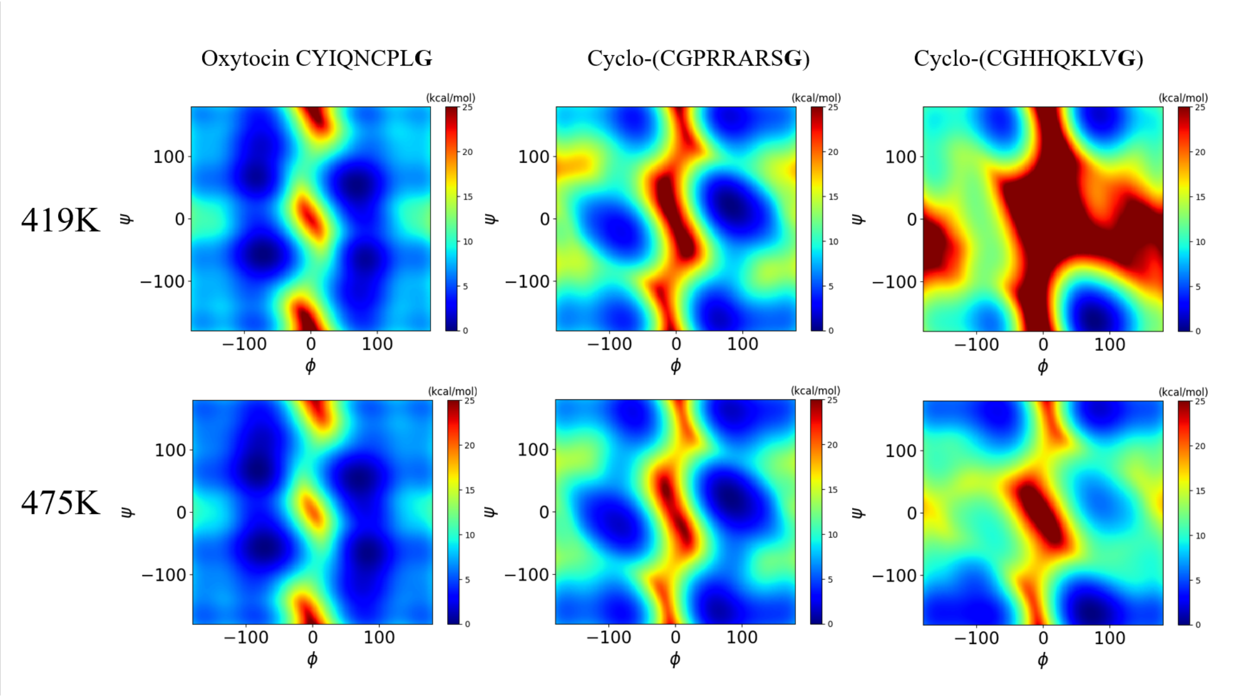


Figure S2: The free energy surface along the $G_{\phi}{}^{9}$ and $G_{\psi}{}^{9}$ Ramachandran space, for all the conventional MD ensembles of the three cyclic peptides examined in the main text, at temperatures 419K and 475K. A bolded font is used for the glycines being analyzed.

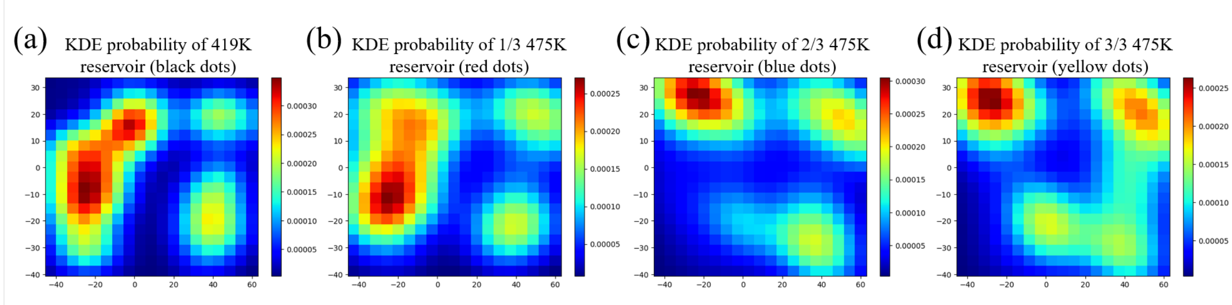


Figure S3: Additional information for Fig. 7. The kernel density estimation probability of reservoir compositions for (a) the 419 reservoir (b) the first 1/3 of the configurations in the 475K reservoir (c) the second 1/3 of the configurations in the 475K reservoir (d) the third 1/3 of the configurations in the 475K reservoir, for cyclo-(CGHHQKLVG).

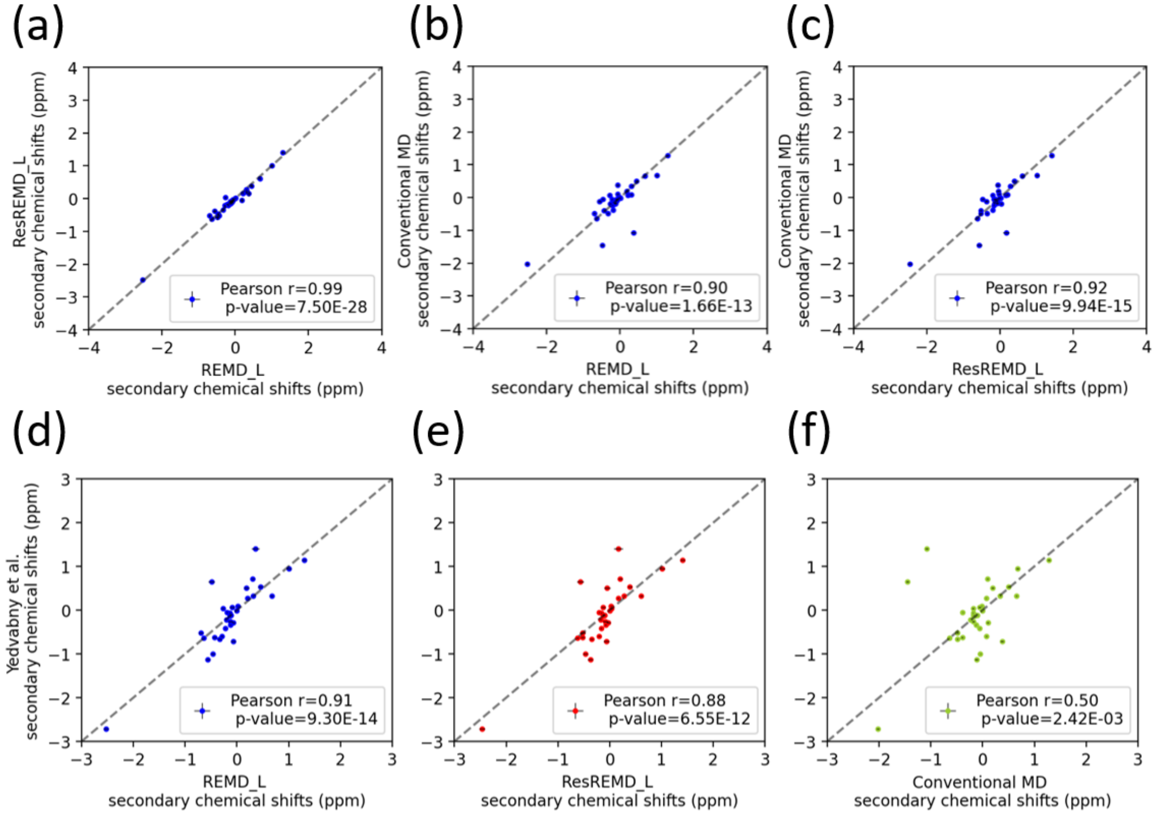


Figure S4: The comparison of the secondary chemical shifts of $C_\alpha$, $C_\beta$, $H_\alpha$, $H_N$ atoms between different pairs of ensembles. (a-c) The mutual correlations of the three computed conformational ensembles (i.e. oxytocin-REMD-L, oxytocin-ResREMD-L, and oxytocin conventional MD) at 300K. (d-f) The three computed conformational ensembles are compared with a past study by Yedvabny et al.[1]

### References

(1) Yedvabny, E.; Nerenberg, P. S.; So, C.; Head-Gordon, T. Disordered structural ensembles of vasopressin and oxytocin and their mutants. *The Journal of Physical Chemistry B* **2015**, *119*, 896–905.
